# Vangl2 regulates the dynamics of Wnt cytonemes in vertebrates

**DOI:** 10.1101/2020.12.23.424142

**Authors:** Lucy Brunt, Gediminas Greicius, Benjamin D Evans, David M Virshup, Kyle CA Wedgwood, Steffen Scholpp

## Abstract

The Wnt signalling network regulates cell proliferation and cell differentiation as well as migration and polarity in development of multicellular organisms. However, it is still unclear how distribution of Wnt ligands is precisely controlled to fulfil all of these different functions. Here, we show that the four-pass transmembrane protein Vangl2 occupies a central role in determining the distribution of Wnt by cytonemes in vertebrate tissue. In zebrafish epiblast cells, mouse intestinal telocytes and human gastric cancer cells, activation of Vangl2 leads to the generation of fewer but extremely long cytonemes, which start to branch and deliver Wnt protein to multiple cells. The Vangl2-activated cytonemes increase Wnt/β-catenin signalling in the surrounding cells. Concordantly, inhibition of Vangl2 function leads to the formation of shorter cytonemes and reduced paracrine Wnt/β-catenin signal activation. A mathematical model simulating the observed Vangl2 functions on cytonemes in zebrafish gastrulation predicts an anterior shift of the morphogenetic signalling gradient, altered tissue patterning, and a loss of the sharpness of tissue domains. We confirmed these predictions during anteroposterior patterning in the zebrafish neural plate. In summary, we show that Vangl2 - a core member of the PCP signalling component - is fundamental to paracrine Wnt/β-catenin signalling by controlling cytoneme behaviour in vertebrate development and tissue homeostasis.

## Introduction

Long-distance cell-cell communication is essential for development and function of multicellular organisms. During embryogenesis, shape-forming signals, called morphogens, are produced at a localized source and act both locally and at a distance to control morphogenesis. In a concentration-dependent manner, morphogens orchestrate the cellular fates in their signalling range by controlling the gene expression of key transcription factors^1-3^. A tightly regulated distribution of morphogens is a prerequisite to allowing their precise spatial and temporal function during embryogenesis in tissue patterning and organ development.

Morphogens of the Wnt signalling family are a class of secreted ligands that can transduce their signals through several distinct pathways to regulate a diverse array of developmental processes^4, 5^. The best-characterized Wnt pathway is the Wnt/β-catenin dependent signalling pathway^6^. Wnt ligands together with Frizzled receptors and the co-receptors Lrp5/6 stabilize the key downstream target β-catenin. The co-transcription factor β-catenin, together with TCF/Lef transcription factors, mediates many cellular processes such as cell differentiation and proliferation and determines tissue patterning along the anteroposterior (AP) body axis. The planar cell polarity (PCP) signalling pathway is a β-catenin-independent pathway in the Wnt signalling network^7, 8^. The Wnt/PCP pathway regulates cytoskeleton remodelling by activation of c-Jun N-terminal kinases (JNK) and members of the Rho-family GTPases such as Cdc42 and RhoA to direct cellular morphogenesis, tissue polarity and cell migration^9^. Within a cell, β-catenin- and PCP-dependent Wnt signalling are well known to act in a mutually repressive manner and inhibiting one will typically upregulate the other.

The transduction of Wnt signalling pathways begins when a Wnt ligand binds to its receptors at the cell membrane. However, the question of how a Wnt moves from a producing cell to fulfil its paracrine function in a tissue remains highly debated^10^. The transport of signal components such as ligands and receptors can be facilitated by signalling filopodia known as cytonemes^11-13^. Indeed, recent high-resolution imaging experiments in zebrafish demonstrated that specialized cytonemes are fundamental in Wnt trafficking in vertebrates^14, 15^. The regulated generation and function of cytonemes is critical as it impacts directly on the signalling range and signalling gradient. Specifically, the number and length of cytonemes generated by a Wnt source cell influence events such as zebrafish neural plate patterning during embryogenesis^14^. However, the molecular mechanism regulating these attributes during zebrafish gastrulation to allow neural plate patterning is still unclear.

Unlike in flies, Wnt-dependent activation of the receptor-tyrosine kinase-like orphan receptor 2 (Ror2) is thought to act as a crucial receptor of the Wnt/PCP pathway^16^, and in turn drives *de novo* biogenesis of Wnt8a-positive cytonemes^17^. The subsequent formation of cytonemes is influenced by activation of cytoskeletal regulators such as the small Rho GTPase Cdc42, which controls actin polymerisation^14, 18^. Ror2 is thought to function by forming a protein complex with other PCP regulatory proteins at the plasma membrane, at a sub-membrane region, or at cell-cell junctions. This PCP core complex can include the key component, the four-pass trans-membrane protein, Van-Gogh-like (Vangl) 1/2 in vertebrates. As Vangl2 lacks any known receptor or enzymatic activity^19^, proteinprotein interaction domains of Vangl2 are likely to modulate downstream signalling. Upon Wnt activation, the kinase domain of Ror2 phosphorylates Vangl2 to activate downstream signalling, such as c-Jun N-terminal kinase (JNK) signalling^16, 20^. Supporting a functional role for Vangl2, its knockdown inhibits axon outgrowth by inhibition of filopodia formation^21^. It has been suggested that this downstream signalling pathway is triggered in a Wnt concentration-dependent manner in mouse limb development^22^. During zebrafish gastrulation, Vangl2 is asymmetrically localised at the plasma membrane and localizes to growing filopodia^23^. It is currently unclear if the Vangl2-positive filopodia perform signalling functions in tissue organization.

Here we show that Vangl2 - together with Wnt8a and Ror2 - is loaded on cytoneme tips. Vangl2 positive cytoneme tip complex activates JNK signalling to increase cytoneme length, enhance cytoneme stability and thus the number cytoneme contacts and their contact time. We find that Vangl2-mediated cytoneme stabilization is vital for paracrine Wnt/β-catenin signalling in both human tissue culture and zebrafish embryo. Concordantly, blockage of Vangl2 function or JNK signalling leads to a quick collapse of signalling filopodia. Consequently, impairment leads to a reduction of both Wnt dissemination and paracrine Wnt signalling in human cancer tissue culture, the zebrafish embryo and the mouse intestinal crypt. Based on our *in vitro* findings, we developed a mathematical model of how changes in cytoneme length, stability, and contacts would affect embryogenesis. This mathematical model of morphogen distribution in the zebrafish gastrula predicts that increased Wnt cytoneme stability and contact time leads to extended signalling range and to altered patterning with fuzzy compartment boundaries. We confirm these predictions *in vivo* during zebrafish neural plate patterning. These findings suggest that the activity of the Vangl2-positive PCP complex determines the emergence of Wnt-positive cytonemes during development and tissue homeostasis in vertebrates.

## Results

### /Vangl2 together with Wnt8a and Ror2 form the cytoneme tip complex

The PCP signalling component Ror2 is an essential regulator of Wnt8a-positive cytoneme emergence^17^. We investigated the interconnected role of additional PCP family members on cytoneme regulation. The four-transmembrane PCP protein Vangl2 is a PCP core member that is localised at the tips of filopodia extending from both neurons as well as in forming membrane protrusions in gastrula cells during zebrafish gastrulation^21, 24, 25^. Vangl2 has been suggested to be activated via phosphorylation by Ror2 kinase in mouse^22^.

To analyse the localisation and function of PCP components together with Wnt8a, we first sought to establish an *in-vitro* test system to monitor cytoneme behaviour using zebrafish PAC2 fibroblast cells in culture. First, we asked if PAC2 cells transported endogenous Wnt8a by filopodial protrusions. We found that these fibroblasts have numerous, dynamic filopodial protrusions and that endogenous Wnt8a can be detected on these filopodia, similar to over-expressed fluorescently-tagged Wnt8a (Fig. 1A, Supplementary Figure 1A,B). Therefore, we define filopodial protrusions as Wnt8a-bearing cytonemes when they co-localized with fluorescently-tagged Wnt8a along the filopodium or at the filopodium tip (Fig. 1A,B). We measured fluorescent intensity along the PAC2 cytonemes (Fig. 1G) and found that glycophosphatidylinositol (GPI)-anchored, membrane-bound mCherry (mem-mCherry) was evenly distributed along the length of cytonemes from tip to base, whilst Wnt8a-GFP was predominantly localised to the tip of the cytoneme (Fig. 1B,H). Next, we transfected plasmids encoding Ror2-mCherry and GFP-Vangl2 to investigate the localization of these PCP proteins in the PAC2 cells. Ectopically expressed Ror2-mCherry was membrane localized (Fig. 1C) with a slight increase in fluorescence of Wnt8a intensity at the tip (Fig. 1I). GFP-Vangl2 and Ror2-mCherry expression was membrane localized and present all along the cytonemes (Fig. 1D,J), whereas GFP-Vangl2 can be also seen at the filopodia tip of PAC2 cells (Supplementary Fig 1C). GFP-Vangl2 accumulated dynamically together with Wnt8a at cytoneme tips (Fig. 1E) with an increase in fluorescence intensity at the tip (Fig. 1K). We find that in the presence of over-expressed untagged Vangl2, Ror2-mCherry accumulated more strongly at the cytoneme tip (Fig.1 F, L). This shows that Vangl2 is present on cytonemes and accumulates with other cytoneme complex members such as Wnt8a and Ror2 at the tip of the cytoneme, suggesting a potential role in emergence and function of Wnt cytoneme.

**Figure 1:**
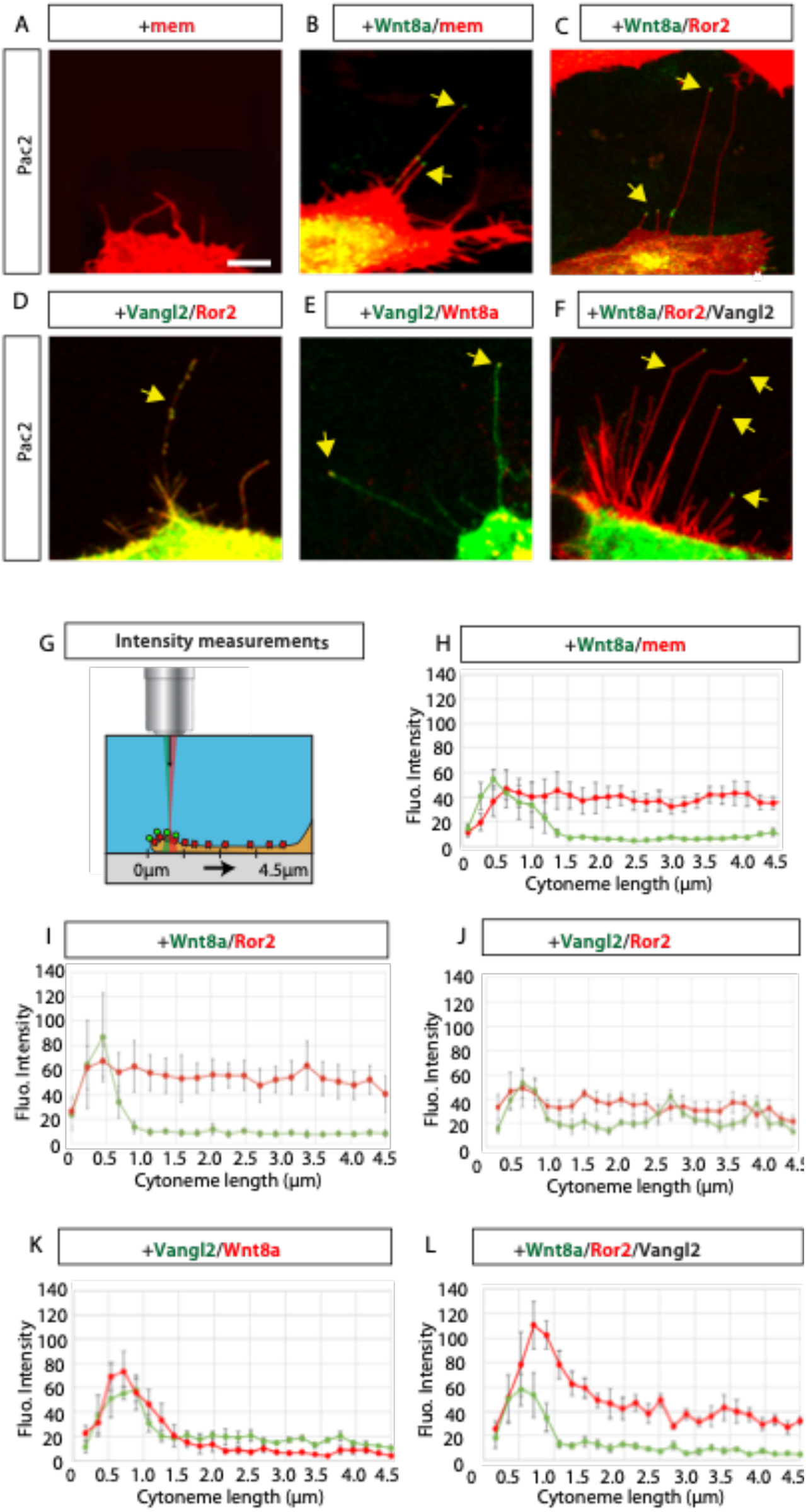
Vangl2 is present on the tips of Wnt8a positive cytoneme. PAC2 zebrafish fibroblasts transfected with indicated constructs and analysed 24h post transfection. Yellow arrows indicate Wnt8a on tips of cytonemes. White scale bar equals 5μm. (G): Fluorescent intensity measurements along cytoneme length from tip at 0μm to 4.5μm along cytoneme. (H-L): Fluorescent intensity analysis (Grey Value) of tagged proteins Wnt8a-GFP, Wnt8a-mCherry, Mem-mCherry, Ror2-mCherry, and GFP-Vangl2, relative pixel intensity values were measured along cytoneme length starting at the cytoneme tip, N=5 filopodia. Standard error of the mean (SEM)=1.

### Vangl2 stabilizes cytonemes in zebrafish fibroblasts

To characterise the function of Vangl2 in cytoneme formation, we analysed the effect of Vangl2 on cytoneme number, length, and stability. PAC2 fibroblasts have numerous, dynamic filopodial protrusions, which are positive for Wnt8a. We found on average 5.7 cytonemes per cell in control cells with an average length of 7.9 μm (Fig. 2A,G,H, Supplementary Fig. 2A,B). The average cytoneme length was slightly longer than Wnt8a-negative filopodia lengths (Supplementary Fig.3,4). We found that 23% of the observed cytonemes were 10 μm or longer (Fig. 2I). Transfection of Ror2 significantly increases the number of filopodia, however, the length and number of cytonemes are unaltered (Fig. 2B, G, H, Supplementary Fig. 3B,G,H). Activation of Vangl2 caused a 58.5% reduction of cytonemes per cell, however, the average cytoneme length increased significantly by 187.3% (Fig. 2C,H, Supplementary Fig. 2B). While in control cells only 23% of cytonemes were > 10μm, in the Vangl2-overexpressing cells 68.9% were > 10 μm, and 43.8% were > 20 μm (Fig 2I). Ror2/Vangl2 double-transfected cells had a similar and significant increase in Wnt8a positive cytoneme length (Fig. 2D,H, Supplementary Fig. 2B), and in addition displayed more branching (Fig. 2D). While the length of cytonemes increased, the average number of cytonemes per cell in Vangl2 and Vangl2/Ror2 cells was reduced by 58.5% and 41.3%, respectively (Fig. 2G, Supplementary Fig. 2A). Phospho-mutants of Vangl2 exhibit a dominant negative effect on signalling (Gao et al., 2011), and so we used an N-terminal deletion mutant of Vangl2 (ΔN-Vangl2) lacking the two Ser/Thr phosphorylation clusters to block Vangl2-mediated signalling. Consistently, ectopic expression of ΔN-Vangl2 led to a reduction of in the number of long cytonemes and an increase in the number of short cytonemes (Fig. 2E,F,H,I Supplementary Fig. 2A,B). Co-expression of Ror2 did not reverse the phenotype (Fig. 2F, G, H), suggesting that Vangl2 functions downstream of Ror2.

**Figure 2:**
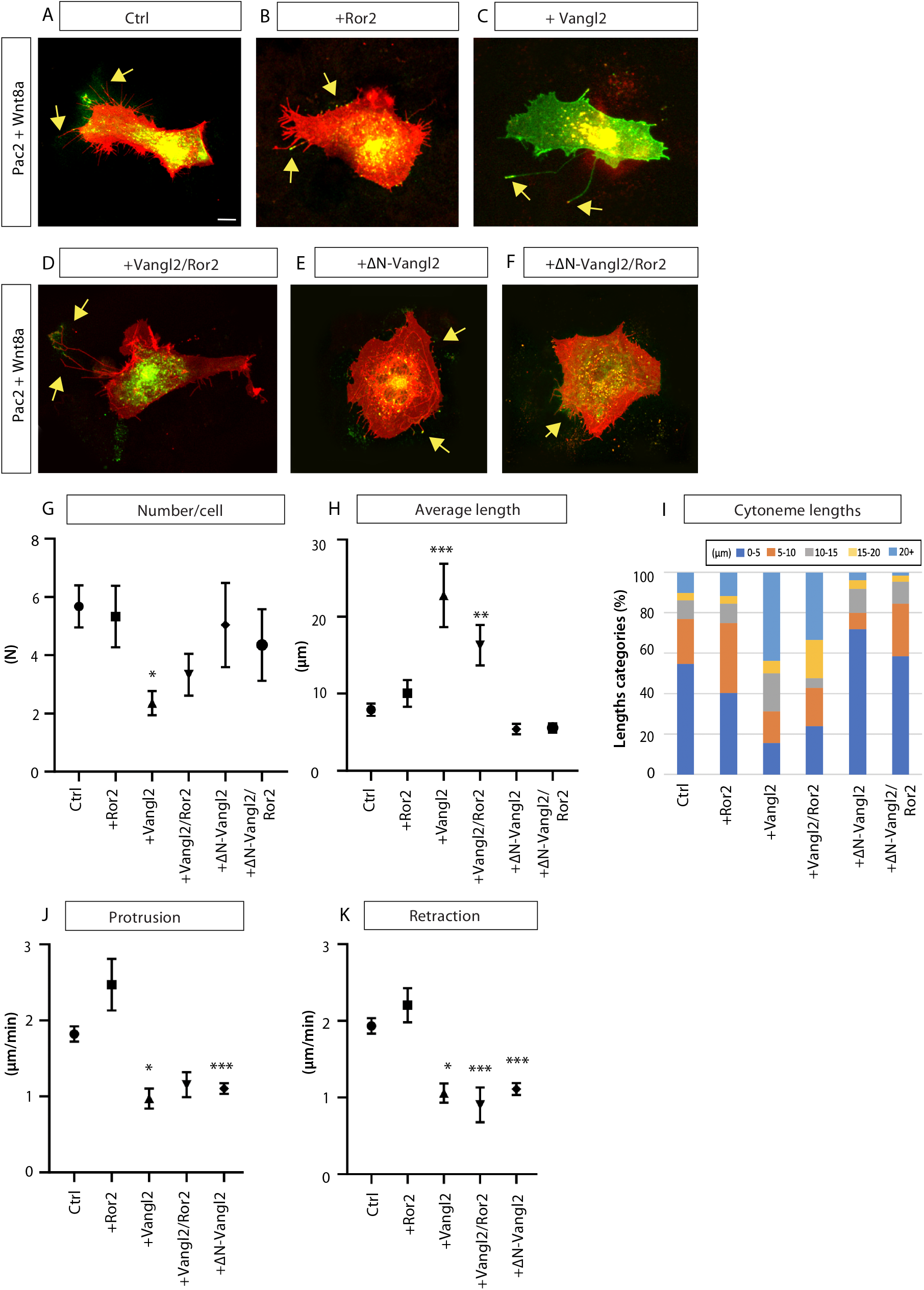
Vangl2 controls the emergence of Wnt8a cytonemes in fibroblasts. (A-F): Wnt8a-positive PAC2 zebrafish fibroblasts transfected with indicated constructs, imaged and analysed 24h post transfection. Yellow arrows indicate examples of Wnt8a positive cytonemes. Scale bar= 10μm. (G): Number of Wnt8a positive cytonemes per cell (N = number). (n= 25, 9, 14, 6, 25, 14 cells). (H): Length of Wnt8a positive cytonemes in PAC2 cells (μm). (n= 139, 52, 32, 21, 131, 65 cytonemes). (I): Breakdown of the percentage of Wnt8a positive cytoneme lengths into 0-5, 5-10, 10-15, 20+μm categories. (J): Protrusion rate (μm/min) of filopodia. (n= 340, 59, 42, 38, 327 timepoints). (K): Retraction rate (μm/min) of filopodia. (n= 340, 59, 52, 41, 327 timepoints). Graphs represent mean and standard error of the mean. Statistical significance: * ≤ 0.05, ** ≤ 0.01, *** ≤ 0.001. (G,H,J,K): Kruskal-Wallis tests with Bonferroni correction for multiple tests. SEM=1. Corresponding dot plots are shown in Supplementary Figure 2, and analysis of PAC2 filopodia is shown in Supplementary Figure 3 & 4.

Next, we asked how Vangl2 increases the length of cytonemes. In Vangl2 and Vangl2/Ror2 transfected cells, many long protrusions appear stabilized and less dynamic compared to cytonemes of control cells or cells transfected with Ror2. This is reflected in a significant reduction in both protrusion and retraction rates by 46.6% and 45.2%, respectively when Vangl2 is transfected (Figure 2J,K, Supplementary 2C,D). The reduced protraction and protrusion rates reflect long periods where cytonemes are firmly attached to neighbouring cells. ΔN-Vangl2 transfected cells also exhibit a significant reduction in retraction rate, possibly due to immobility of short protrusions in these cells or that the N-terminal of Vangl2 is not required for Vangl2 function in cytoneme dynamics. These data suggest that Vangl2 plays a role in cytoneme regulation, specifically involved in stabilizing cytonemes and, therefore, increasing cytoneme contact times.

### Vangl2 controls the stability of Wnt8a cytonemes during zebrafish gastrulation

We next investigated the role of Vangl2 in cytoneme regulation *in vivo* in zebrafish embryos. To do this we generated clones of cells by microinjecting mRNA of mem-mCherry and Wnt8a-GFP, together with either Vangl2 or ΔN-Vangl2 mRNA at the 16-cell stage. At 5hpf (50% epiboly), we imaged individual clones of cells in the zebrafish gastrula (Fig. 3A-C). In particular, we focused on visualisation of Wnt8a-positive cytonemes within the embryo. We found that, like what occurred in cultured PAC2 cells, expression of Vangl2 led to significantly fewer but longer cytonemes per cell compared to control (Fig. 3D-F, Supplementary Fig. 5A,B). The average length of cytonemes significantly increased of 39.6% (Fig. 3E,F). Expression of ΔN-Vangl2 did not significantly change the number or the length of cytonemes compared to control (Fig. 3C,D-F), suggesting that the observed lengthening requires Vangl2 function. The cytonemes in Vangl2-expressing embryonic cells were likewise found to branch abnormally (Fig. 3B, white asterisk), form multiple contact points and extend over larger areas between cells compared to control (Fig. 3I, J; Supplementary Figure 5E).

**Figure 3:**
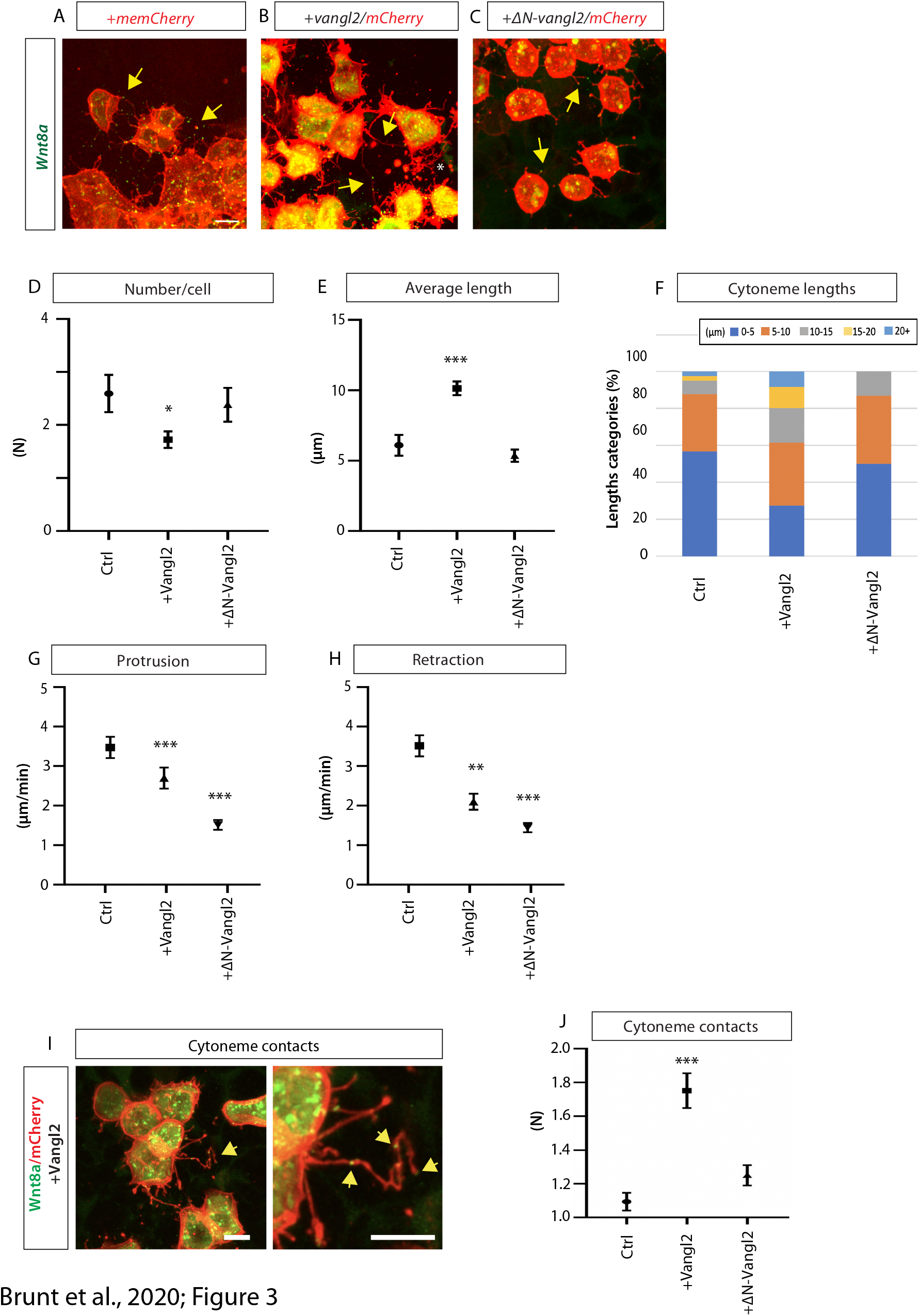
Vangl2 regulates length and stability of Wnt8a cytonemes in the zebrafish embryo. (A-C): Zebrafish embryos were injected with indicated constructs and imaged at 5hpf. Yellow arrows show examples of Wnt8a positive cytonemes. Scale bar= 10μm. (D): Number of Wnt8a positive cytonemes per cell (N = number). (n= 3, 6, 3 embryos, n= 27, 123, 21 cells). (E): Length of Wnt8a positive cytonemes (μm). (n= 81, 269, 54 cytonemes). (F): Breakdown of the percentage of Wnt8a positive cytoneme lengths into 0-5, 5-10, 10-15, 20+μm categories. (G): Protrusion rate (μm/min) of filopodia. (n= 3, 5, 3 embryos, n= 70, 50, 104 timepoints). (H): Retraction rate (μm/min) of filopodia. (n= 3, 5, 3 embryos, n= 86, 54, 104 timepoints). (I): Vangl2, Wnt8-GFP and mem-mCh expressing cells in the 50% epiboly embryo exhibiting cytonemes with multiple Wnt8a contact points. (n= 32, 125, 80). Scale bar= 10μm. (J): Quantification of number of Wnt8a-positive contact points per cytoneme. Graphs represent mean and standard error of the mean. Corresponding dot plots are shown in Supplementary Figure 5. Statistical significance: * ≤ 0.05, ** ≤ 0.01, *** ≤ 0.001. (D,E,G,H,J): Kruskal-Wallis tests with Bonferroni correction for multiple tests. SEM=1.

Cytonemes on cells in the control embryos were very dynamic and protruded and extended at approximately 3.7 μm/min, while in the Vangl2-expressing cells, cytonemes were more static, with increased contact time (Fig. 3G-H, Supplementary Fig. 5C,D). This was due to a significant reduction in both protrusion of 22.7% and retraction rates of 40.6% compared to control. Addition of ΔN-Vangl2 also significantly reduced cytoneme protrusion and retraction rate compared to control (Fig. 3C, G, H). Therefore, we conclude that Vangl2 regulates Wnt8a cytoneme length and protrusion dynamics by allowing tips to anchor - both *in vitro* and *in vivo*.

### Vangl2-mediated JNK activation is required to stabilize cytonemes

We next wanted to investigate the mechanisms by which Vangl2 regulates cytoneme stability. Wnt/PCP can transduce signals through the receptors Frizzled and Ror2 to Rac-JNK and RhoA-ROCK signalling cascades in a context-dependent manner^26^. Apart from its nuclear functions, JNK also directly regulates the cytoskeleton by phosphorylation of diverse cytoplasmic targets including proteins directly interacting with the actin cytoskeleton, as well as microtubule-associated proteins^27^. Similarly, the RhoA-ROCK signalling cascade can induce actin cytoskeletal re-organization and cell movement^7^. In addition, there is crosstalk between these pathways; RhoA can activate JNK during convergent extension movement in Xenopus, and loss of RhoA can be rescued by over-expression of JNK1^28^.

We recently reported results from a cell-culture-based screen to identify kinases that regulate cytoneme formation^17^. In this screen, in addition to Ror2, we identified several key family members of the JNK signalling pathway including MKK4 and JNK3 as positive regulators of Wnt8a cytoneme length. After receiving external signals, MAP kinase kinases (MKK) phosphorylate and activate c-Jun N-terminal kinases (JNK). In turn, the JNKs phosphorylate a number of transcription factors, primarily components of AP-1^29^. In our screen, forced expression of MKK4 led to an increase in the average length of cytonemes by 36.4%, whereas JNK3 expression resulted in an increase of 19.7%. To further probe the involvement of JNK signalling, we carried out a reporter assay in HEK293T cells, which express a very low level of endogenous Wnt ligands and a defined set of Fzd receptors^30^. In cells with low JNK activity, the KTR-mCherry reporter^31, 32^, is localised to the nucleus (Fig. 4A). However, upon activation of JNK signalling, phosphorylation of the JNK-KTR-mCherry reporter causes it to translocate to the cytoplasm^31^. Control HEK293T cells have low JNK activity, shown by nuclear localisation of KTR-mCherry (Fig. 4B,C). Transfection of Ror2, Vangl2 or a combination of both did not significantly change the ratio of cytoplasmic to nuclear signal (Fig 4Bii-iv,C). Remarkably, addition of either Wnt5a or Wnt8a protein led to a significant increase in JNK activity in Ror2/Vangl2-expressing cells, as seen movement of KTR-mCherry from nucleus to cytoplasm (Fig. 4Bv-vi,C). This suggests that Wnt protein with Ror2 and Vangl2 initiates JNK signalling in HEK293T cells. Wnt5a or Wnt8a did not activate JNK signaling in cells transfected with ΔN-Vangl2 (Fig. 4Bvii-viii,C). This indicates that Vangl2 is a key downstream element in Ror2-Wnt activated JNK signalling in HEK293T cells.

**Figure 4:**
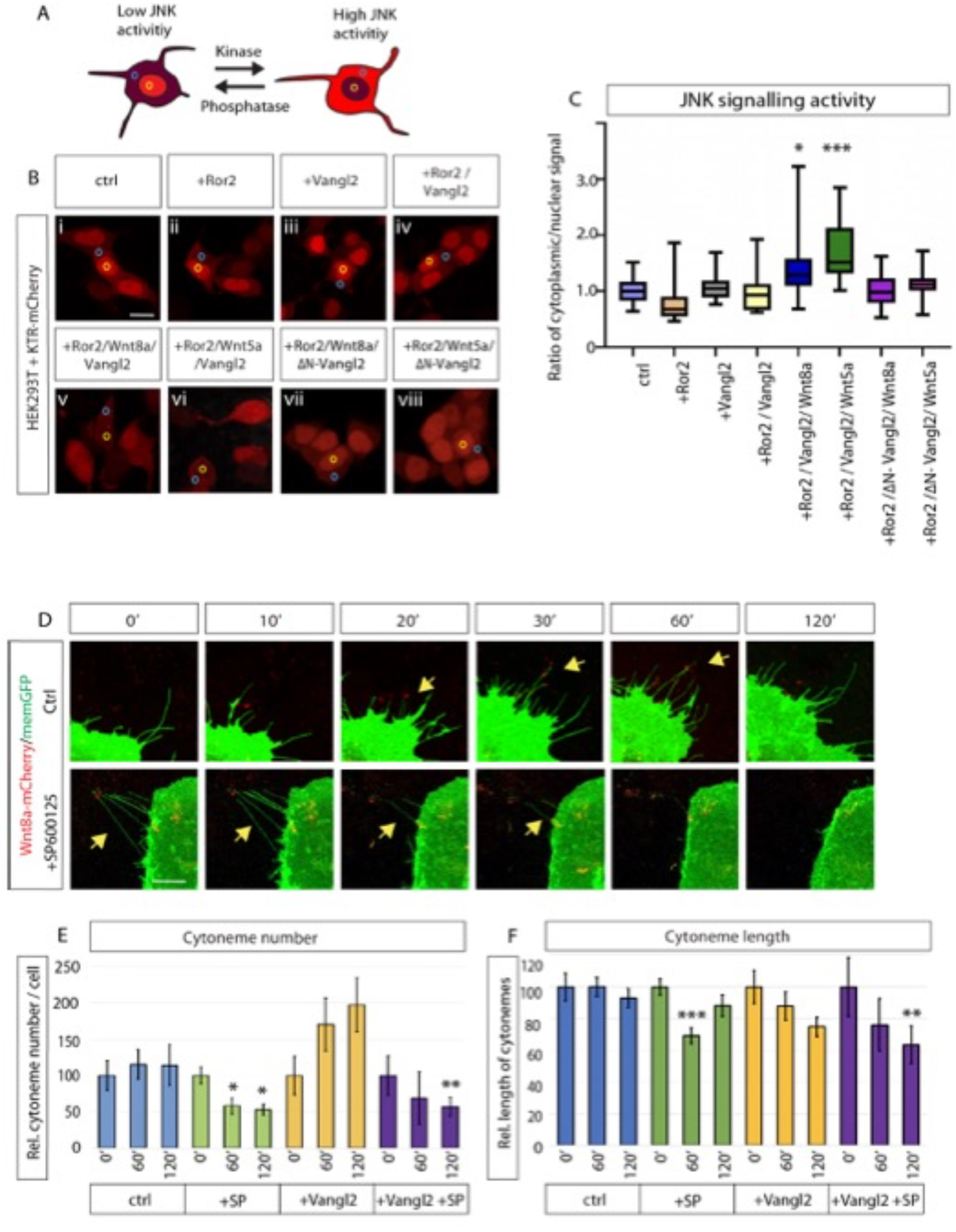
Vangl2 activates JNK signalling to control the formation of Wnt8a cytonemes. (A): Schematic of KTR-mCherry JNK reporter in HEK293T cells. Low JNK activity in the cell leads to nuclear localisation of KTR-mCherry (yellow circle). High JNK activity leads to phosphorylation and a switch in KTR-mCherry localisation to the cytoplasm and a reduction in nuclear localisation (blue circle). (B): HEK293T cells all transfected with KTR-mCherry plus Ror2 (ii); Vangl2 (iii); Ror2 & Vangl2 (iv-vi); Ror2 & ΔN-Vangl2 (vii-viii). Wnt8a protein and Wnt5a protein (500ng/ml) was also added 24hrs prior to imaging (v-viii). (C): Box and whisker plot of normalised ratio of cytoplasmic to nuclear signal of KTR-mCherry. Increased ratio indicates increasing JNK activity. (n= 27, 25, 30, 24, 28, 29, 49, 60 cells). Chart shows median and whiskers show 99^th^ percentile. Kruskal-Wallis tests with Bonferroni correction. SEM=1. (D): Time series of PAC2 cells at 0 min, 10 min, 20min, 30min, 60min and 120min in control cells and cells post 20μM SP600125 treatment. (E): Relative number of filopodia per cell in relation to time=0hrs, at 0min, 60min, 120min. (n= 3, 6, 10, 4 cells. n= filopodia at 0hr, 1hr, 2hr = (102/111/109, 251/156/122, 158/185/215, 58/50/44). Stars over +Sp600125 60’ & 120’ (green bars) significant to 0’. Stars over Vangl2+SP600125 120’ (purple) significant to Vangl2 120’ (yellow). (F): Relative filopodia length (μm) in relation to time=0hrs, at 0min, 60min, 120min. (n= 3, 6, 10, 4 cells. n= filopodia at 0hr, 1hr, 2hr = (102/111/109, 251/156/122, 158/185/215, 58/50/44). Star over +Sp600125 60’ (green) significant to 0’. Stars over Vangl2+SP600125 120’ (purple) significant to Vangl2 0’ (yellow). Statistical significance: * ≤ 0.05, ** ≤ 0.01, *** ≤ 0.001. (E-F): Kruskal-Wallis tests without Bonferroni correction. SEM=1. Scale bar= 10μm.

We observed that Wnt8a cytonemes form and retract within some tens of minutes (Fig. 2 & Fig. 3 & ^14^). To assess the role of JNK signalling in this process, we treated Wnt8a-GFP/mem-mCherry transfected PAC2 cells with a small molecule inhibitor of JNK kinase activity. In vertebrates SP600125 specifically inhibits all three JNKs (JNK1-3) within minutes without inhibition of ERK1 or −2, phosphop38, or ATF2^33^. We then recorded and analysed the effect of JNK blockage on protrusion length and number (Fig. 4D-F). We found that Wnt cytonemes collapse and retracted following the addition of SP600125 (Fig. 4D), whereas cytonemes in DMSO treated PAC2 cells are unchanged. A time course revealed that over the course of 120 mins, control cells showed no significant change in the number or length of signalling filopodia (Fig. 4E,F). However, JNK inhibition caused a significant reduction in average relative protrusion length and number after 1 hour. Vangl2-expressing cells, which had longer protrusions (as in Fig 2H) prior to the addition of SP600125, also had a significant change in the number and length of cytonemes compared to untreated Vangl2-expressing fibroblasts. A detailed analysis of cytoneme lengths showed a specific loss of extremely long signalling filopodia as a result of JNK inhibition (Supplementary Fig. 6A). We also tested the involvement of RhoA/ROCK signalling as it can be similarly activated downstream of PCP signalling to regulate cytoskeletal re-arrangement. To do so, we treated cytoneme-bearing cells with Y-27632, an antagonist of Rho-associated kinase (RhoA/ROCK)^34^. While JNK inhibition had a significant effect at 60 min, Y-27632 treatment produced only a modest and non-significant reduction of protrusion length after 5 hrs (Supplementary Fig. 6B-F). We conclude that JNK signalling is the primary signalling cascade required for fast cytoneme formation downstream of Wnt/Vangl2/Ror2 signalling and it contributes to the formation and stabilisation of long Vangl2-positive cytonemes.

### Vangl2-controlled cytonemes regulate Wnt/β-catenin signalling in neighbouring cells

Next, we investigated the effect of Vangl2 on cytoneme-mediated Wnt protein delivery by investigating Wnt/β-catenin signal activation in the receiving cells (Fig. 5). To measure paracrine signal activation, we used a sensitive reporter system of AGS gastric cancer cells - which are primed for Wnt/β-catenin signalling - transiently transfected with a Wnt/β-catenin reporter with seven TCF-responsive elements driving expression of nuclear mCherry (7xTCF-nls-mCherry^35^). AGS cells have also been shown to exhibit Ror2-dependent Wnt8a cytonemes^17^. We co-cultivated AGS cells transiently expressing combinations of Wnt8a, Vangl2, Ror2 and ΔN-Vangl2 together with the STF-mCherry reporter cells (Fig. 5A,B). The expression of nuclear mCherry significantly increased in the receiving cells when Wnt8a was expressed in the source cells (Fig. 5Bii,C). This suggests that Wnt8a can activate the Wnt/β-catenin pathway in a paracrine way. Expression of Vangl2 alone in the source cells did not induce a Wnt/β-catenin response in the receiving cells (Fig. 5Biii,C). However, Wnt8a/Vangl2 cotransfection, which we showed in zebrafish cells increased cytoneme numbers and length, could significantly increase STF reporter activation in receiving cells in comparison to control and Wnt8a only (Fig. 5Biv,C). The significant increase of Wnt signal transmission to the responding cells is dependent on Vangl2, because co-transfection of Wnt8a/ΔN-Vangl2 showed no detectable effect on reporter activation compared to Wnt8a expressing source cells (Fig. 5Bv,C). Next, we tested if the Vangl2-dependent increase of paracrine Wnt/β-catenin signal activation depends on Ror2 function. We found that expression of Wnt8a/Ror2/Vangl2 in the producing cells further increased reporter activation in the receiving cells (Fig. 5Bvi,C), and this was abrogated by co-expression of with ΔN-Vangl2 or kinase-dead Ror2 (Ror2^3i^) (Fig. 5Bvii,Bviii,C). These data indicate that the activity of Wnt8a in source cells is markedly enhanced by two factors known to increase cytoneme number and length, consistent with the transmission of the Wnt8a signal by cytonemes.

**Figure 5:**
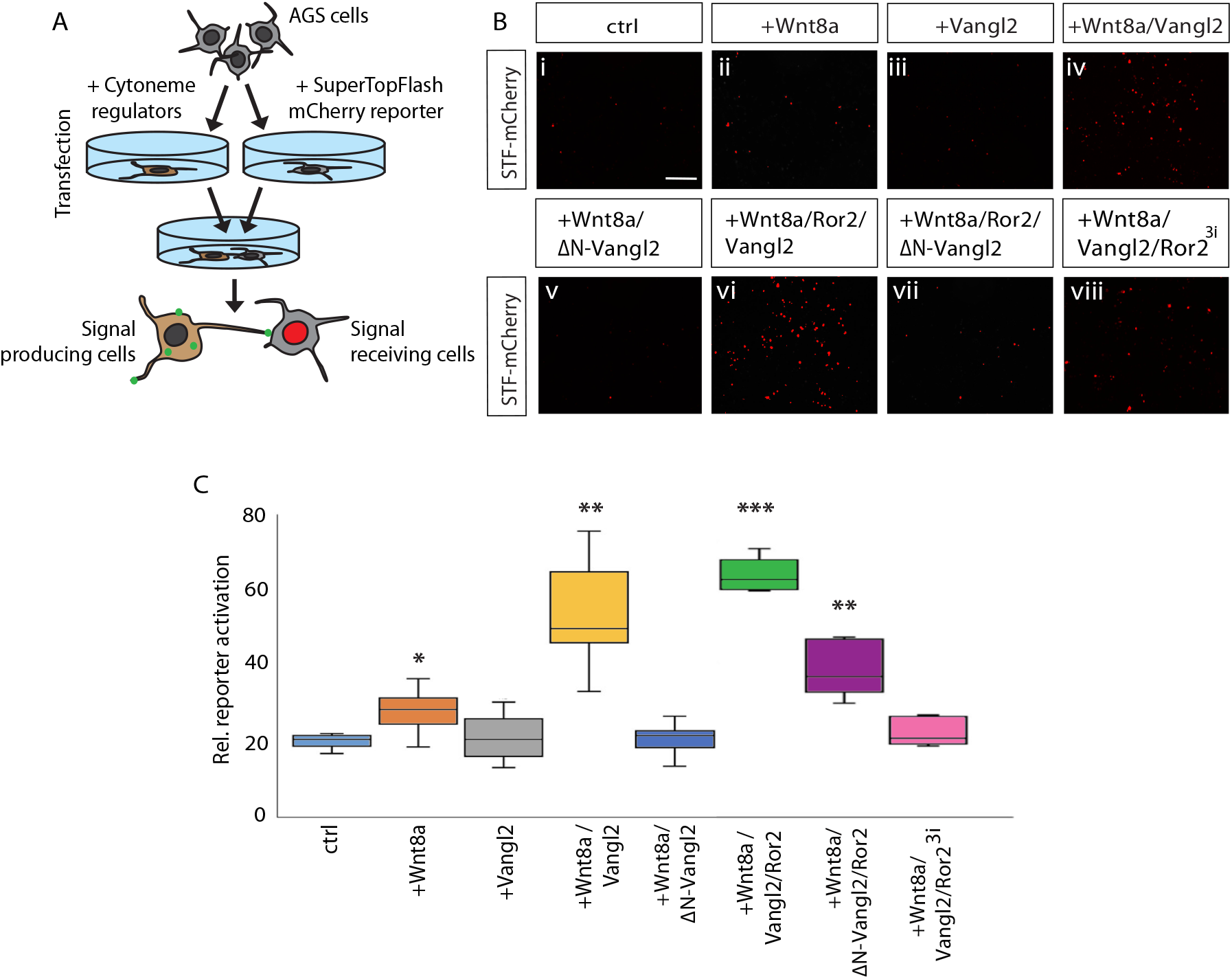
Vangl2 activity in the Wnt source cells regulates paracrine Wnt/β-catenin signal activation. (A): Schematic of SuperTOP Flash (STF) reporter co-cultivation assay in AGS cells. AGS cells transfected with STF reporter were co-cultivated AGS cells transfected with combinations of with Wnt8a, Vangl2, Ror2 and ΔN-Vangl2 plasmid. (B): AGS cell STF reporter activation at each condition (i-viii). Scale bar= 10μm. (C): Relative STF reporter activation in cells when co-cultured with control; Wnt8a; Vangl2; Wnt8a/Vangl2; Wnt8a/ΔN-Vangl2, Wnt8a/Ror2/Vangl2; Wnt8a/Ror2/ΔN-Vangl2 and Wnt8a/Vangl2/Ror2^31^. (n= 5,10,5,5,5,5,5,5 repeats). Data represented as box and whisker plots. Oneway ANOVA tests. Statistical significance: * ≤ 0.05, ** ≤ 0.01, *** ≤ 0.001.

To test for the requirement for cytoneme-mediated Wnt transport in a further vertebrate tissue, we investigated the influence of Vangl2 on Wnt8a cytonemes in the mouse intestinal crypt. The stem cells at the bottom of intestinal crypt require a constant supply of Wnt signalling for tissue homeostasis^36-39^. *In vivo*, subepithelial myofibroblasts, described as PDGFRα+ telocytes, are the major source of physiologically relevant Wnts to maintain these intestinal crypts^40-42^. The ability of these myofibroblasts to provide essential Wnts is compromised when they lack cytonemes as a result of siRNA mediated ROR2 inhibition^17^. We used crypt organoids as a system to test whether telocytes also require Vangl to generate cytonemes to distribute Wnt proteins in the mouse intestinal crypt. Vangl1 and Vangl2 expression in telocytes was reduced by siRNA-mediated knockdown (Fig. 6A). The number of filopodia was significantly reduced after Vangl1 knockdown and even more markedly reduced after double knockdown of *Vangl1/Vangl2* (Fig. 6B,C). In parallel, we found a reduction in filopodia length in all three knockdown experiments with the most significant reduction after double knockdown of Vangl1/Vangl2 (Fig. 6D). Next, we used an organoid formation assay to analyse the requirement for Vangl-dependent Wnt cytonemes (Fig. 6E). Organoids of Wnt-deficient *Porcn^-/-^* crypt cells need to be co-cultivated with WT telocytes for maintenance (Fig. 6E,F). Telocytes extend filopodia to contact crypt organoids and supply them with signalling factors such as Wnt proteins. After simultaneous knockdown of Vangl1 and Vangl2 in the Wnt-producing telocytes, we observed a strong decrease in the number of organoids. This suggests that the Wnt cytonemes from the telocytes are required for the induction and maintenance of the intestinal crypt organoids and that Vangl1/2 are crucial for their formation.

**Figure 6:**
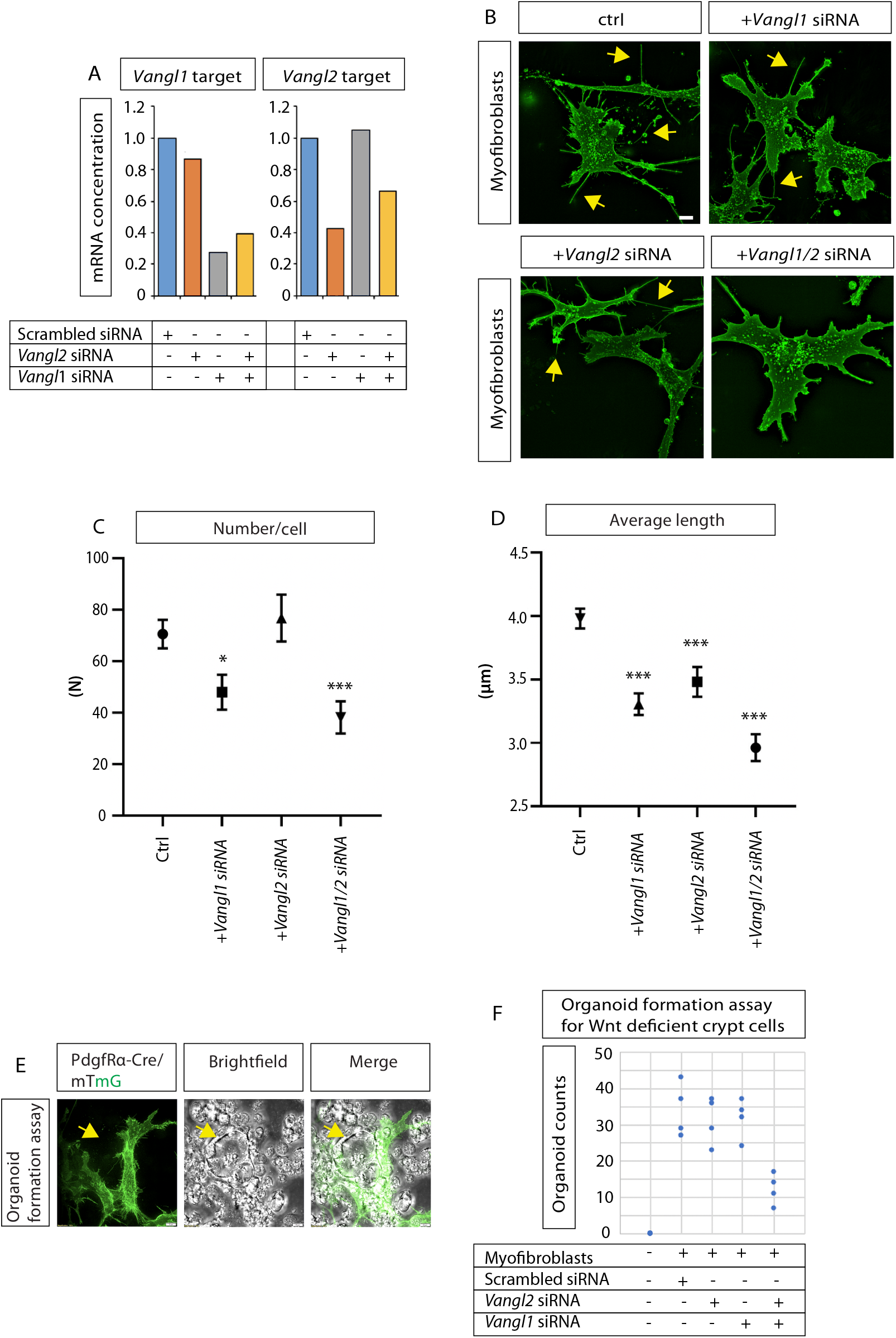
Vangl function in murine telocytes is required for the formation of Wnt deficient intestinal crypt. (A): mRNA concentration of Vangl1 and Vangl2 in scrambled siRNA, Vangl1, Vangl2 and Vangl1/Vangl2 siRNA treated murine telocytes. (B): Telocytes stained with FITC-phalloidin. Yellow arrows indicate filopodia protrusions. Scale bar= 10μm. (C,D): Number of filopodia per cell and average length of filopodia in telocytes treated with control scrambled siRNA and siRNA for Vangl1, Vangl2 and Vangl1/Vangl2. C: (n=35, 21, 22, 27 cells). D: (n= 61, 50, 30, 30 cells, n= 3488, 1796, 1964, 1079 filopodia). Graphs represent mean and SEM. Statistical significance: * ≤ 0.05, ** ≤ 0.01, *** ≤ 0.001. (C,D): Kruskal-Wallis tests with Bonferroni correction for multiple tests. SEM=1. (E): Organoid formation assay: PDGFRα-cre/Rosa2-mTmG telocytes (green), co-cultured with Porcn^-/-^-intestinal epithelial cells. Both cell types visualised in brightfield and merge to show organoid formation. Yellow arrows indicate telocyte filopodia. (F): Organoid formation assay of Porcn deficient crypt cells cocultured with murine telocytes treated with control scrambled, Vangl1, Vangl2 or Vangl1/Vangl2 siRNA. Organoid counts were recorded per condition. (n= 4 assays per condition).

### Simulation predicts an important role for Vangl2-controlled cytonemes in the zebrafish gastrula

On the basis of these findings, we hypothesized that Vangl2 function in the Wnt source cells is crucial for Wnt dissemination via cytonemes which we examined *in silico*. To quantitatively test the consequences of altered Vangl2 function on gradient formation and tissue patterning in the zebrafish neural plate, we created an agent-based simulation of morphogen distribution via cytonemes using the Chaste modelling software^43, 44^. First, we generated a 2D model of the zebrafish gastrula based on the positional information of every cell during the first 10hrs of zebrafish gastrulation^45, 46^, representing a portion of the overall gastrula. We defined the population of marginal cells as Wnt8a source cells and the overlying epiblast cells as Wnt-receiving neural plate cells (Fig. 7A,B). The simulation takes into account ligand transport by cytonemes, ligand decay and the migration and proliferation of epiblast cells using the agent-based simulation approach (Fig. 7C-F). We employed cytonemes as the exclusive transport mechanism from the producing marginal source cell group to the target cell group. We made the assumption that all cells are initially at the animal pole then migrate and intercalate to produce a thin tissue, which covers the yolk during the epiboly movement and Wnt transport process. We found that cytonemes can distribute Wnt8a in a graded manner in the dynamically evolving target tissue of the zebrafish embryo during gastrulation. Cells receiving a high concentration of Wnt8a acquire hindbrain fate, according to a pre-defined Wnt threshold, whereas, cells receiving a lower concentration acquire forebrain/midbrain fate. We further find the formation of a stable boundary between midbrain and hindbrain (MHB) due to a sharp drop of the morphogen concentration across the boundary. Next, we tested two scenarios with varying ligand concentration and varying cytoneme number, stability and growth properties (to match experimentally observed properties), based on our in vivo measurements after alteration of Vangl2 function (Fig. 3). We found that increasing ligand concentration (in these simulations by a factor of 10) within the morphogenetic field leads to an anterior shift of MHB in comparison to the control situation (Fig. 7G,H). We found that increasing lengths but reducing the mean number of cytonemes per cell after expression of Vangl2 in the source cells (by the experimentally determined factor of 33%) leads to a slight anterior shift of the MHB (Fig. 7I). Remarkably, co-expression of Vangl2 and Wnt8a leads to a significant broadening of the hindbrain territory (Fig. 7J). Notably, we also found the MHB becomes less distinct, (Figure 7B), whereby there are considerably more cells exhibiting a fate incongruous with that defined by their position relative to the computed boundary. On the basis of the simulations, we predicted a strong increase in the range of Wnt signal activation as well as a loss of robust boundaries in the neighbouring tissue when Vangl2 function is accelerated in the Wnt8a source cells.

**Figure 7:**
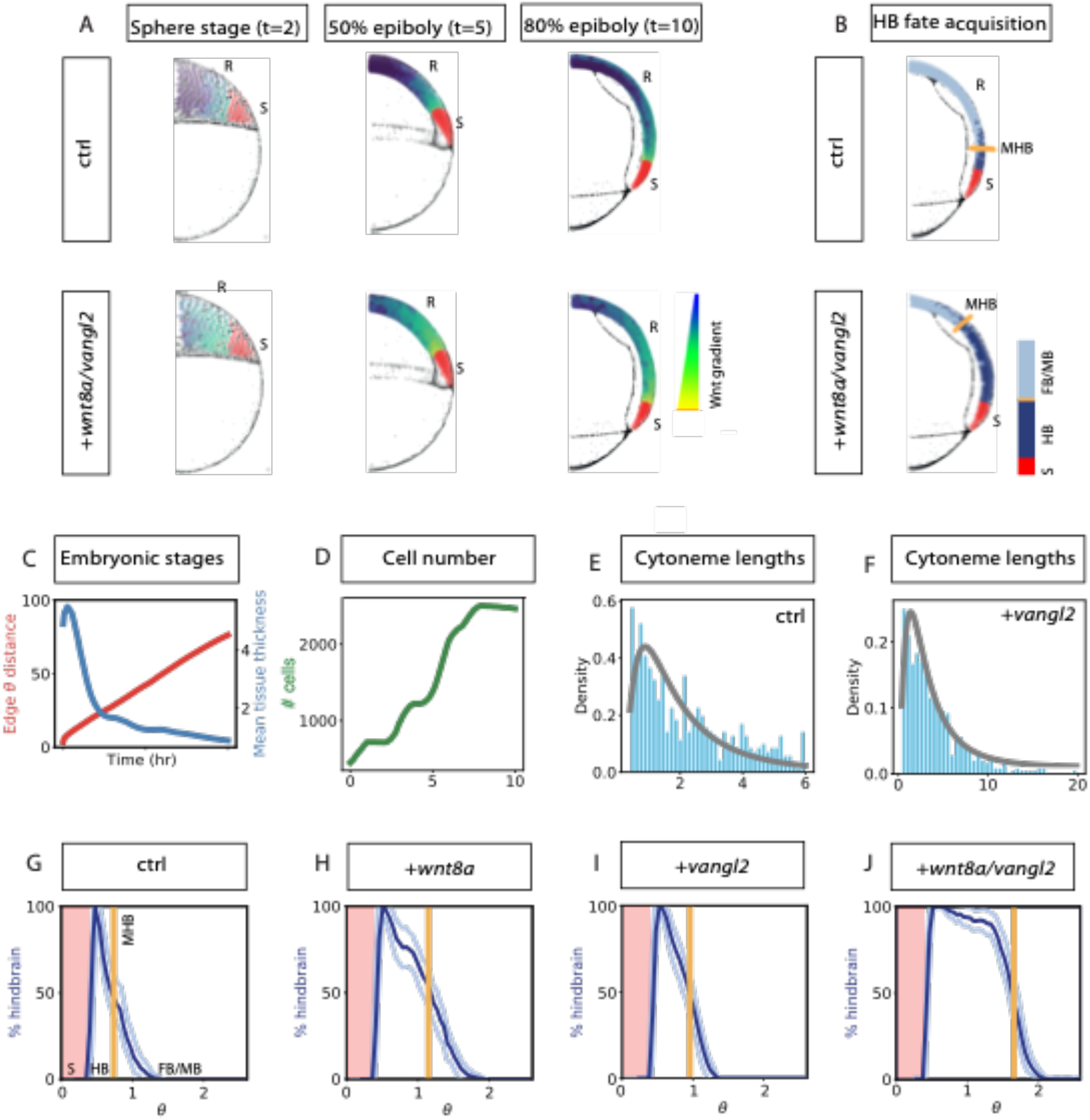
Agent-based modelling predicts an essential role for Vangl2 in Wnt-mediated tissue patterning during zebrafish gastrulation. (A-B): Comparison of Wnt protein distribution in control zebrafish embryos with zebrafish embryos with Vangl2/Wnt8a overexpression. State of one realisation of the model at the indicated time point (50% epiboly=5hpf; 80%epiboly=8.5hpf) for base parameter values and Wnt8a and Vangl2 over-expression parameters. For the 80% epiboly panel, the fate of the cells is also plotted in (B). Source cells are indicated in red. The colours of the cells in anterior tissue correspond to the relative level of Wnt8 protein received (A). For ease of viewing, Wnt8a protein values have been normalised by the maximum value attained by any cell across the simulation and log transformed. In the cell fate diagram (B), cells acquiring a hindbrain fate are marked in dark blue, whilst those not acquiring a hindbrain fate are marked in grey. The orange line marks the estimated midbrain-hindbrain boundary (MHB). (C): Tissue growth properties - a single simulation. Note that the mechanics of the tissue growth are preserved across all conditions and so this graph is representative of all simulations. The red curve shows the proportion of the yolk that is covered by the tissue at the indicated times. The blue curve shows the mean tissue thickness (in terms of the number of cells). (D): Evolution of the cell number. (E-F): Histograms of cytoneme lengths at the final state of the simulation (normalised by cell diameter) for the control parameters (E) and Vangl2 over-expression parameters (F). The grey curve shows a fit of the data to a log-normal distribution with means 2.0 (for E) and 3.8 (for F). (G-J): Distributions of cell fates over the angular polar coordinate at the end state of the simulation according to a hindbrain Wnt8a threshold of 100 (AU) for control parameters (G), Wnt8a overexpression parameters (H), Vangl2 overexpression parameters (I) and Wnt8a / Vangl2 overexpression parameters (J). The dark and light blue curves respectively show the mean and standard deviations of the proportion of cell fates acquiring a hindbrain fate in 100 equi-spaced bins around the yolk over 100 model simulations for each condition. The orange line shows the estimated position of the MHB. The red shaded area marks the position of the margin of wnt8a producing cells.

### Vangl2 activity in the Wnt source cells regulates neural plate patterning in zebrafish neurogenesis

To test our prediction from the simulation, we analysed the consequences of Vangl2 function on the Wnt signalling range in zebrafish embryogenesis. In detail, we wanted to understand if the changes to cytoneme length and stability, as a result of Vangl2 alterations, impacted on neural plate patterning as suggested by agent-based simulation. First, we analysed the formation of the Wnt source tissue. The zebrafish embryonic margin functions as a major signalling source for Wnt and Fgf in the early gastrula. The transcription factor Notail (ntl; ortholog in mouse TBXT, formerly known as T or brachyury) is essential for the induction of signalling factors such as Wnt8a and Fgf8a at the embryonic margin^47^. *Ntl* expression is detected early in development in Wnt8a positive mesodermal progenitor cells and is required for body axis formation. Wnt8a and Ntl act in a positive autoregulatory loop to reinforce their expression^48^. Therefore, we analysed the expression pattern of *ntl* at the embryonic margin after altering of expression of Wnt8a and Vangl2. At 60% epiboly, the overexpression of Vangl2; and more drastically, Wnt8a with Vangl2, caused an abnormal broadening and ectopic expression pattern of *ntl* (Fig. 8A-D).

**Figure 8:**
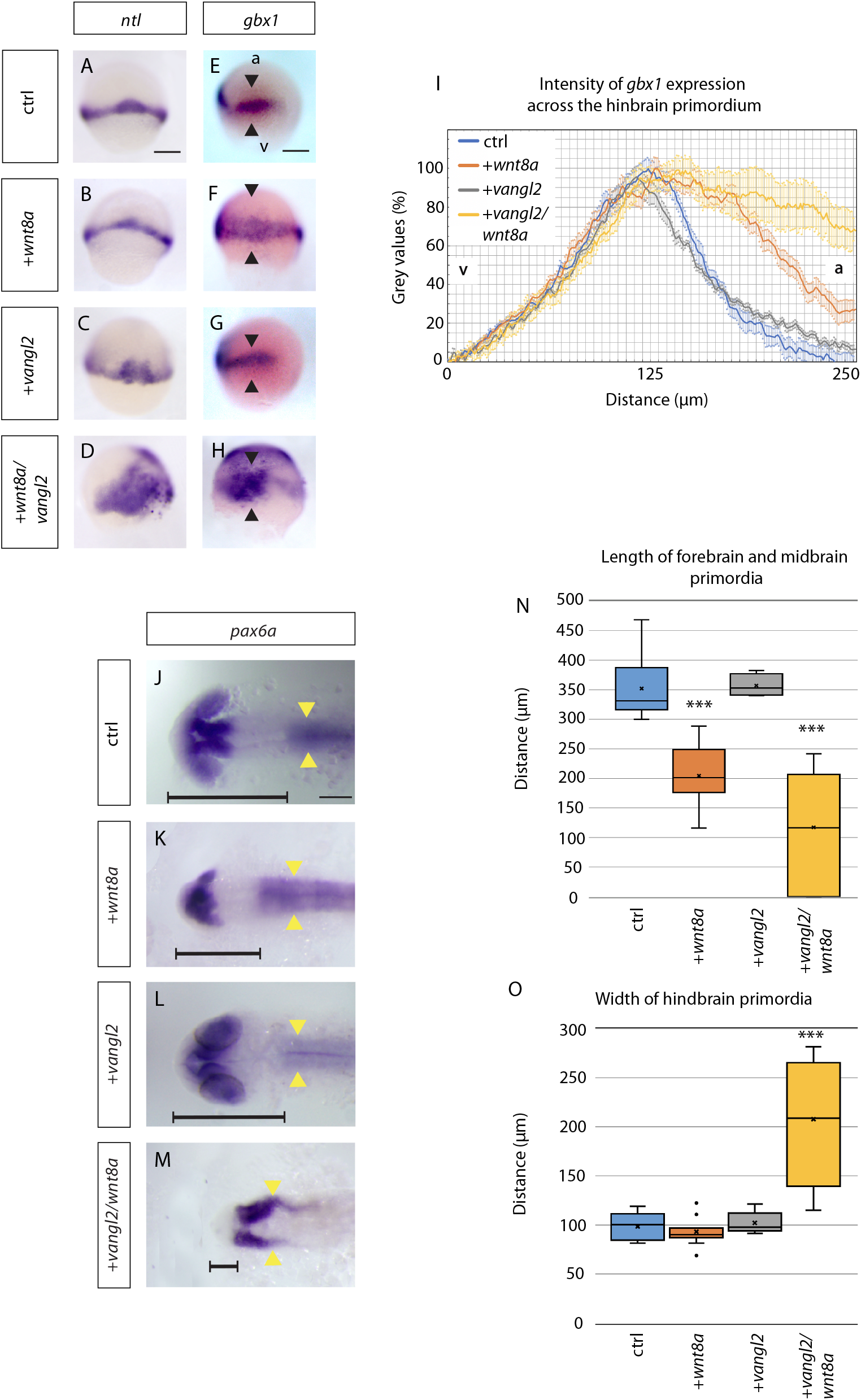
Vangl2 function is crucial for anteroposterior patterning of the zebrafish neural plate. (A-H): *in situ* hybridisation analysis of indicated markers in 60% epiboly zebrafish embryos (6.5hpf) injected with Wnt8a, Vangl2 and Wnt8a/Vangl2. Scale bar indicates 200μm. (*ntl*: n= 35, 16, 6, 12 embryos) (*gbx1*: n= 6, 19, 3, 11 embryos). a= animal pole, v=vegetal pole. Black arrowheads indicate width of *gbx1* expression. (I): Intensity of *gbx1* expression from *in situ* hybridisation (grey values in %) across the hindbrain primordium from vegetal (v) to animal (a) pole in embryos injected with mRNA for the indicated constructs at 60% epiboly (6.5hpf): Control-blue, *wnt8*-orange, *vangl2* - grey, *wnt8a/vangl2*-yellow. (J-M): *in situ* hybridisation to mark *pax6a* expression in the primordia of the forebrain and the hindbrain in embryos injected with mRNAs for the indicated constructs at 24hpf: Control, Wnt8a, Vangl2 and Wnt8a/Vangl2. Scale bar indicates 100μm. Horizontal black line indicates length of forebrain and midbrain primordium. Yellow arrowheads show the width of the hindbrain primordium, indicating the extent of convergent and extension of these cells. (n= 13, 19, 8, 10 embryos). (N): Box and whisker plot of length of forebrain and midbrain primordia in control (blue), Wnt8a (orange), Vangl2 (grey) and Wnt8a/Vangl2 (yellow) 24hpf larvae (μm). Measured from anterior forebrain *pax6a* expression to the position of the midbrain-hindbrain boundary (MHB) shown by horizontal black line (J-M). (n= 6, 12, 4, 10 embryos). (O): Box and whisker plot of maximum width of hindbrain primordia in control (blue), Wnt8a (orange), Vangl2 (grey) and Wnt8a/Vangl2 (yellow) 24hpf larvae (μm). Measured from maximum width of hindbrain *pax6a* expression shown by yellow arrowheads (J-M).. (n= 5, 11, 5, 8 embryos). Statistical significance: * ≤ 0.05, ** ≤ 0.01, *** ≤ 0.001. (N,O): ANOVA tests. SEM=1.

Next, we mapped the expression pattern of neural plate markers. *Gbx1* is a further direct Wnt signalling target in the gastrulating embryo and a marker for hindbrain identity^49^. *gbx1* mRNA expression can be detected at the marginal region at 7hpf (60-80% epiboly) (Fig. 8E-H). By measuring the intensity of the *gbx1* expression from vegetal to animal pole, we were able to measure the width of the expression pattern, as well as the sharpness of the MHB (Fig. 8I). Wnt8a expression leads to a broadening of the expression pattern of *gbx1* compared to control and a reduction in the sharpness of the MHB (Fig. 8F,I). Vangl2 alone has little effect on *gbx1* expression, with similar expression distance and sharpness of boundaries to control. However, Wnt8a and Vangl2 overexpression together leads to a more exaggerated broadening to the *gbx1* expression domain pattern compared to control and Wnt8a alone. There is a reduced sharpness of the midbrain-hindbrain boundary (MHB), as a result of more ectopic *gbx1* expression further from the original expression domain. As predicted from our simulation, in synergy, Wnt8a and Vangl2 leads to a broadening and less sharp *gbx1* expression domain, suggesting that the increase in cytoneme length and stability may affect the signalling range and capability to signal to further cells. This has an impact on the specification of the brain primordia, for example, the *gbx1* positive hindbrain primordium.

By 24hpf, primordial brain boundaries have formed due to activation of specific markers for brain regions. *Pax6a* is expressed at the forebrain primordium and at the hindbrain primordium and rhombomeres. Addition of Wnt8a posteriorizes the brain, leading to reduction of forebrain and midbrain primordia structures (Fig. 8J-N). Vangl2 overexpression leads to similar anteroposterior patterning as control. However, Wnt8a plus Vangl2 leads to significantly reduced forebrain and midbrain structures, with complete abolishment of forebrain primordia structures in some cases. This is more severe than Wnt8a alone, though not significantly so. Emergence of convergent extension (CE) phenotypes with addition of PCP Vangl2 is only evident after the addition of Wnt8a (Fig. 8J-M,O). Therefore, as well as a CE phenotype, Wnt8a and Vangl2 causes a neural plate patterning phenotype, with reduction of forebrain and midbrain primordia structures. Overall, Vangl2 alterations leads to changes in early neural plate patterning, as outlined by changes to *ntl* and *gbx1* expression. At later stages, CE and AP patterning phenotypes of the neural tube are present. We suggest that the observed effect on AP patterning of the zebrafish brain anlage is a result of increased Wnt cytoneme length and cytoneme contact times, impacting on an increased signalling activation range.

## Discussion

The PCP signalling pathway was initially characterized in *Drosophila* and orchestrates cell polarity across an epithelium^50^. For example, in Drosophila, PCP is responsible for the coordinated and consistent orientation of hairs and bristles. In vertebrates, PCP homologues regulate the orientation of inner ear sensory hair cells, hair follicles of the skin, and epithelial cells bearing multiple motile cilia. PCP is further involved in regulating cell migration, differential adhesion across cells, orientation of cytoskeletal elements, and positioning of cell extensions, such as filopodia^7^. Core members of the Wnt/PCP signalling in vertebrates are, for example, Vangl and Ror2^51^. The transmembrane factor Ror2 serves as a Wnt co-receptor helping to relay the signal to Vangl. After activation by the Wnt5a ligand, the receptor tyrosine kinase Ror2 phosphorylates two Ser/Thr phosphorylation clusters in the cytoplasmic N-terminal region of the Vangl2 protein^22^, with subsequent Dvl2 phosphorylation^52^. In mouse, mutants for the core PCP component Vangl2 exhibit open neural tubes (craniorachischisis)^19, 53^. Similarly, human mutations in both Vangl1/2 are associated with spina bifida^54, 55^. In zebrafish, the *Vangl2* mutant *trilobite* exhibits a broadened body axis, owing to similar defects in convergent extension (CE) movements during development^56^. Therefore, Vangl2 together with Ror2 take centre stage in the PCP signalling pathway.

### Vangl2 and cytoneme formation

Here, we demonstrate that Vangl2 has an essential function during formation of Wnt cytonemes and thus in the distribution of Wnt ligands across vertebrate tissues. The mechanism of cytoneme-based ligand transport has been observed in many tissues^11, 15, 57^. Initially, cytonemes have been described in various tissues of Drosophila transporting Fgf, Dpp and HH signalling components^58-61^. Recent data suggest that cytonemes are also used to mobilize signalling components in vertebrates. Shh is transported by cytonemes in the chick limb bud and uses a Cdc42-independent mechanism^62^. Cytonemes are also fundamental for transporting Wnt signals. We have shown that cytonemal transport of Wnt8a is essential during neural plate patterning during zebrafish gastrulation^14^. Wnt is loaded on cytonemes and can be found at the cytonemal tip (Fig. 1). In chick, there is evidence that also the Wnt receptor Frizzled7 (Fzd7) is required for somite formation and Fzd7 puncta could be detected on cytonemes emitting from the ectodermal layer^63^. Activation of Wnt signalling in stem cells can be similarly mediated via Wnt receptor positive cytonemes^64^. These findings are similar to observations in Drosophila, in which cytonemes containing Fzd receptors extend from myofibroblasts to pick up Wg signal from the wing disc^65^. However, the molecular mechanism underlying cytoneme formation is still unclear.

Here, we show that Vangl2 can be seen together with Wnt8a and Ror2 on cytoneme tips of PAC2 fibroblasts as well zebrafish epiblast cells *in vivo*. These data are supported by observations in zebrafish epiblast cells and zebrafish hindbrain motor neurons, in which Vangl2 localizes to filopodia tips^24, 25^. Similarly, in the mouse neural tube, Vangl2 and Frizzled3 are enriched on the tips of growth cone filopodia^21^. However, the function of Vangl2 in cytonemes is yet to be elucidated.

We provide evidence that Vangl2 regulates the appearance of Wnt-bearing cytonemes: Vangl2 activation induces specifically long, stable and branched cytonemes, which carry Wnt protein at their tips. Besides its function in cell polarity and coordinated cell migration, Vangl has been suggested to play an important role in cellular protrusion formation. In Drosophila, knock-down of vangl and prickle lead to the formation of very few and short Fgf cytonemes^66^. In Vangl2^-/-^ / *Loop-tail* mouse mutants, filopodia were unable to extend^67^. In hippocampal neurons, Vangl2 was found to regulate dendritic branching, with Vangl2 knockdown leading to reduced spine density and dendritic branching^68^. In zebrafish, Vangl2 modulates the formation and polarization of actin-rich, large protrusions in ectodermal cells^25^. Vangl2 is enriched at membrane domains that are developing these large protrusions compared with non-protrusive domains. Interestingly, Vangl2 has been suggested to destabilize protrusions, whereas Fzd3a is required to stabilize the same extensions^24^. However, the nature of these protrusions is unclear. Here, we investigated the function of Vangl2 on cytonemes - small, slender protrusions, which form and retract within minutes and are loaded with signalling proteins. We show that Vangl2 function is essential for the formation of long Wnt cytonemes.

### PCP/Vangl2 signalling regulates JNK signalling

The mechanism of how Vangl2 regulates cytoneme appearance is unknown. Our data suggests that cytoneme emergence can be modulated by an intrinsic signalling cascade and Rac/Jun N-terminal kinase (JNK) and Rho-associated kinase (ROK) participate in the Wnt-mediated PCP pathway. The Rac/JNK signalling cascade is an intracellular relay pathway and is essential in regulating both the cytoskeleton and cell adhesiveness^69^. At the core of this cascade are the stress-activated MAP kinase kinases MKK4 and MKK7 that activate JNK to modulate cytoskeletal and nuclear events^70, 71^. During dorsal closure in Drosophila, JNK signalling regulates the formation of actin and myosin dependent protrusions^72, 73^. We have previously identified several members of the Rac/JNK signalling family as regulators of Wnt8a positive cytonemes^17^. We showed that overexpression of MKK4 and JNK3 led to the formation of longer Wnt8a cytonemes in PAC2 cells. Furthermore, we have shown that blockage of Rac1 (and Cdc42) mediated signalling by ML141 leads to a collapse of Wnt cytonemes within 2h^14^. Here, we show that Vangl2-positive stable cytonemes are retracted within minutes after JNK signal inhibition (Fig. 4).

Indeed, there is evidence of Vangl2 mediated regulation of JNK activity. Vangl2/PCP signalling leads to activation of JNK and c-Jun by phosphorylation^70, 74^. Furthermore, Vangl2 regulates cell adhesion by regulating Rac1/JNK activity at adherens junctions^75^. Similarly, it has been suggested that Vangl2 promotes phosphorylation of c-Jun and AP-1 in zebrafish^76^. It has been further suggested that p62 is required to recruit and activate JNK through an evolutionarily conserved VANGL2-p62-JNK signalling cascade in Xenopus and human breast cancer cells^77^. We conclude that Vangl2/PCP activates Rac1/JNK signalling to regulate cytoneme generation in the Wnt source cell.

In addition to JNK signalling, Wnt/Vangl stimulation can also induce activation of RhoA-dependent signalling. RhoA/Rock regulate cell migration^78^ - specifically convergence and extension (CE) movements - by mediating Wnt/PCP signalling in zebrafish and Xenopus gastrula^28, 79-81^. Although there are reports of cross-regulation and synergism, RhoA/Rock signalling has been rather implicated in changes in cell shape, orientation, and polarity whereas Rac/JNK signalling is involved in filopodia formation^82^. This notion is supported by our observation, that JNK signalling is essential for controlling the formation of highly dynamic cytonemes, whereas RhoA/ROCK signalling seems to be more important for adjusting the number of filopodia^83^ or cytonemes^14^ presumably as an adaption to an altered cell morphology. Indeed, we found that Cdc42 can regulate the number of cytonemes^14^ similar to RhoA, which regulates cell shape and protrusion lifespan in CE movements^82^. Therefore, we hypothesize that Vangl2 activates Rac/JNK signalling to regulate the elongation and collapse of Wnt cytonemes within minutes and, therefore, influences the dynamic distribution of Wnt ligands on cytonemes.

In addition to the intrinsic function of Vangl2 on Rac/JNK, it has been suggested that Vangl/PCP signalling influences protrusion formation by influencing the composition of the extracellular matrix in Drosophila and zebrafish^25, 66^. In the zebrafish Vangl2 mutant *trilobite*, longer and thicker protrusions were observed. However, it is unclear if these extensions are filopodia, which are used for signalling or nanotubes forming intracellular bridges^84^. In Drosophila, it has been suggested that the PCP components prickle and vangl are essential for Fgf cytonemes formation. After blockage of prickle / vangl function, cytonemes were reduced in number and length when *pkRNAi* or *VangRNAi* were expressed^66^. In these flies, the composition of the extracellular matrix (ECM) was altered, suggesting that prickle and vangl are involved in maintaining normal levels of glypicans such as Dally, Dlp and laminin. However, a molecular mechanism explaining the relation between PCP signalling and ECM composition is unclear.

### Vangl2 cytonemes and paracrine signalling and tissue patterning

Recently, we could show that Wnt distribution by cytonemes is essential during a very narrow time window of 2hr to achieve neural plate patterning during early zebrafish gastrulation^85^. This raised the question of how to reach a sufficiently high flux of Wnt protein into the field by fast cytonemes, given the fact that only a few cytonemes form per Wnt producing cell. Here we show that Vangl2 activation is a crucial regulator of cytoneme behaviour regarding stability, length, and growth properties, and the ability to branch out with the potential to contact multiple target cells. This observation became relevant for the generation of the morphogenetic field in our simulation. When the simulation assumed that all cytonemes deliver the ligand to a single neighbouring cell and we take into account the actual number of cytonemes, we observed only minimal differences in the concentration of the ligand within the morphogenetic field. However, as soon as we allow multiple contact sites per cytoneme over a longer time, we increase the level of activation in the target cells combined with a wider patterning activation (Fig. 7). We concluded that for generating a signalling gradient employing the mechanism we describe it is essential to regulate all parameters of cytonemes. In particular, the net change in the flux of Wnt8a from producing to receiving tissue controls the range of the gradient and is directly proportional to the filopodia number and length but more importantly, also to the number of contact events per cytoneme. We observed that activation of Wnt cytoneme transport by Vangl2 and Ror2 led to significant upregulation of Wnt/β-catenin signalling in the neighbouring cells, whereas blockage of Ror2 or Vangl2 function led to a strong reduction of paracrine Wnt signalling. Consequently, activation of Vangl2 and Wnt8a led to a synergistic upregulation of paracrine Wnt/β-catenin signalling and in posteriorization of the neural tube during zebrafish gastrulation. In future work, the predictions of our model could be validated by investigating paracrine signal activation and tissue patterning. We argue Vangl2 function is essential to control stability and multiple contact events given the short time window in which Wnt signalling is required for neural plate patterning.

## Conclusion

Cytonemes are taking centre stage in cell-cell communication in invertebrates^57^ and there are an increasing number of examples similarly describing the essential function of these signalling filopodia in vertebrates^15^. However, our understanding of how these cytonemes are regulated is still in its infancy. Here, we describe Vangl2/PCP-induced cytonemes as transport carriers for Wnt8a in zebrafish. In cell culture experiments, we use PAC2 fibroblasts and HEK293T cells to provide further evidence for the importance of Vangl2-dependent regulation of cytonemes in Wnt trafficking. In addition, we show that human gastric cancer cells AGS process paracrine Wnt signalling via cytonemes, which are influenced by Vangl2 activity. Then, we use murine intestinal stroma cells, which express multiple Wnts to maintain the Wnt gradient operating in the intestinal crypt and provide evidence that also these Wnts utilize Vangl1/2-dependent cytonemes for their transport. Finally, we show that Vangl2 activity controls neural plate patterning in the developing zebrafish embryo by controlling Wnt8a cytonemes. In summary, we show that autocrine PCP pathway activation via Vangl2 induces Wnt cytonemes in the Wnt source cell to transmit Wnt to the neighbouring cell to activate paracrine Wnt/β-catenin signalling. We propose that PCP signalling is a crucial mechanism for regulating the emergence of cytonemes to mobilize Wnt ligands in vertebrate development and tissue homeostasis.

## Acknowledgements

Research in the SS lab is supported by the BBSRC (Research Grant, BB/S016295/1 and an Equipment grant, BB/R013764/1) and by the Living Systems Institute, University of Exeter. Studies in the DMV lab are supported by the National Research Foundation of Singapore and National Medical Research Council under its STAR Award Program as well as TIER3 grant MOE2016-T3-1-002. BDE was generously supported by the Wellcome Trust Institutional Strategic Support Award (Grant number 204909/Z/16/Z). KW is supported by MRC Fellowship MR/P01478X/1. We would additionally like to thank Alex Fletcher (University of Sheffield) for assistance with the implementation of the agent-based model and Ned Boulter for programming python wrappers to facilitate parallel processing and analysis of the model simulations. For technical help, we would like to thank Sally Rogers (IF staining), Jordan Kent (STF-assay), and Alexandra Mader (ISH analysis). We would like to thank Cecilia Moens (Fred Hutchinson Cancer Research Center) and Steve Wilson, Masa Tada (UCL) for providing plasmids; Trevor Dale and Toby Phesse (ECSCRI, Cardiff University) for providing the gastric cancer cell lines. Furthermore, we would like to thank Chan Yarn Kit for contributions to Vangl1/2 siRNA work and the entire Scholpp lab for critical comments on the manuscript. We would like to thank the Aquatic Resources Centre (ARC) and the BioImaging Centre, Exeter for excellent technical support.

## Author Contributions

LB and SS designed, performed and analyzed all experiments except where noted and wrote the manuscript. GG and DMV performed the intestinal organoid studies, BDE and KCAW designed and performed the simulations.

## Material and Methods

### Zebrafish Maintenance and Husbandry

WIK wild-type zebrafish (*Danio rerio*) were maintained as previously described at 28°C and on a 14hr light/10hr dark cycle^14^. Experiments were carried out under personal and project licenses granted by the UK Home Office, under ASPA (Animal Scientific Procedures Act), and ethically approved by the Animal Welfare and Ethical Review Body at the University of Exeter.

### Cell Culture

PAC2 zebrafish fibroblasts derived from 24hpf zebrafish embryos were maintained at 28°C in Leibovitz L-15 media (Gibco). PAC2 cells^14^ were kindly provided by Nicholas Foulkes (KIT). HEK293T (Human Embryonic Kidney, CRL-1573) cells from the American Tissue Culture Collection, ATTC, Wesel, Germany were maintained at 37°C with 5% CO_2_ in DMEM media (Gibco). Primary gastric adenocarcinoma (AGS) cells were maintained at 37°C with 5% CO_2_ in RPMI-1640 media (Gibco). All media was supplemented with 10% FBS (Gibco) and 1% Pen/Strep (Gibco) and with L-Glutamine.

### Organoid formation of intestinal crypt cells

Myofibroblasts were prepared from C57BL/6-Tg(PDGFRα-cre)^1Clc/J/RosamTmG^ mice and cultured as previously described^42^. As confluence of cultured cells was reaching 80%, they were transfected with Vangl1 and Vangl2 siRNAs (Dharmacon Cat# J-057276-09-0002 and Cat# J-059396-09-0002 respectively, at 10 nM) using siRNAmax reagent (Invitrogen Cat#13778-030). On day 2 posttransfection, myofibroblasts were mixed with Porcn-deficient intestinal epithelial cells and cultured using RSPO1-supplemented medium. Organoid counting was performed at the time point when the group containing no stromal cells had no surviving organoids left (the end of day 3/beginning of day 4 of co-culture). siRNA-transfected cells were imaged using the Olympus Live Imaging system IX83. Acquired 3D image stacks were de-convoluted using the software cellSens Dimension (Olympus) and are presented as maximum intensity projections.

### Transfection and Plasmids

Cells were transfected using Fugene HD Transfection Reagent (Promega) and imaged after 24-48hr. For the STF reporter co-culture assay, cells were transfected and after 24hr, re-trypsinised to co-culture cells.

Plasmids were used for transfection and to make *in situ* probes and mRNA for microinjection into zebrafish embryos. The following plasmids were used: GPI-anchored mCherry in pCS2+ (Mem-mCh)^86^; zfGap43-GFP in pCS2, xRor2 in pCS2+, XRor2^3i^ in pCS2+ and xRor2-mCherry in PCS2+^17^; zfWnt8a ORF1-mCherry in pCS2+^49^; zfWnt8a ORF1-GFP in pCS2+^14^; meGFP-Vangl2 in pcDNA3.1^24^; zfStbm (Vangl2), in pCS2+, (Addgene, #17067); Stbm-DeltaN-6myc in pCS2+^76^; JNK KTR-mCherry (JNK Kinase Translocation Reporter)^31, 32^; 7xTRE Super TOPFlash-NLS-mCherry (STF-mCherry)^35^.

### Antibody Staining

PAC2 cells were plated on coverslips and transfected with a zfWnt8a-mCherry plasmid using Fugene. 24 hours later, cells were washed once in prewarmed PBS, then fixed using 0.25% glutaraldehyde and 4% PFA at 4°C. Cells were washed in PBS, then blocked and permeabilised using 0.1% Triton X-100, 5% goat serum and 0.2M glycine for 1 hour at room temperature. Cells were washed again, then incubated with anti-Wnt8a antibody MBS9216179 (Generon) overnight at 4°C. Coverslips were washed, incubated with an anti-rabbit secondary polyclonal antibody conjugated to Alexafluor 488 (Abcam 150077) and Phalloidin TRITC for 1 hour at room temperature. Coverslips were washed and mounted using ProLong Mountant (Invitrogen).

### Image acquisition

Cells and zebrafish embryos were imaged on a TCS Leica SP8 confocal microscope, using 63x dip-in objective. Cells were seeded on plastic 35mm dishes and embryos were mounted in 0.7% low-melting point agarose in plastic 35mm dishes and filled with Danieau’s solution.

### Fluorescent intensity along cytoneme

FIJI software was used to plot the pixel intensity along a selected line within an image. A line was assigned from the base to the tip of cytoneme as defined by a membrane marker and the pixel intensity was plotted. The line selection was copied to the ROI manager and the measurement repeated for the second channel.

### Number and length of cytoneme - FIJI Analysis

Membrane protrusions marked with membrane marker were assigned as filopodia. Membrane protrusions marked with membrane marker and positive for tagged Wnt8a were assigned as Wnt8a-positive cytonemes. The number and length of filopodia and cytonemes were measured from base to tip of protrusions in FIJI. Continuous nanotube structures from cell to cell or thicker membrane extensions were discounted.

### Microinjection of mRNA and DNA constructs

Capped sense mRNA was generated from linearized plasmid using the mMessage mMachine SP6 & T7 Transcription Kits (Invitrogen). mRNA was microinjected at the 16-cell stage at 200ng/μl, to generate clonal expression when imaged from 50% epiboly. GFP-Vangl2 DNA and mRNA was microinjected at 2-4 cell for clonal expression for *in situ* experiments.

### KTR-mCherry based JNK reporter assay and analysis

HEK293T cells were transfected with JNK reporter KTR-mCherry and either/or, GFP-Vangl2, xRor2 and Stbm-DeltaN-6myc. Wnt5a and Wnt8a mouse recombinant protein was reconstituted at 100μg/ml in PBS and 0.1% BSA (Biotechne). 500ng/ml Wnt8a and Wnt5a mouse recombinant protein was added and incubated at 37°C for 24hrs before imaging. For analysis, mean grey values were recorded for 3x ROI in the nucleus and 3x ROI in the cytoplasm per cell and an average was taken. The ratio of cytoplasmic/nuclear signal was then recorded and normalized to 1 for the control.

### Inhibitor treatment

Cells were transfected as previously described. 20 μM JNK inhibitor SP600125 (Sigma), dissolved in DMSO, was added to the media. Cells were imaged immediately and 1hr and 2hr after treatment. Y27632 (ROCK inhibitor) was dissolved in water and added to the media at 10 μM for 1, 5 or 24 hrs.

### STF reporter co-culture assay and fluorescent intensity analysis

AGS Cells were transfected with either STF reporter 7xTRE Super TOPFlash-NLS-mCherry or plasmids of interest. After 24hrs, cells were then trypsinised and seeded together for another 24hrs. STF reporter expression was imaged on a Leica widefield microscope. Nuclei STF reporter expression was thresholded in FIJI and relative fluorescence of nuclei was measured.

### Generation of *in situ* probes and *in situ* hybridization

*notail, gbx1* and *pax6a* digoxigenin and FITC antisense probes were generated from linearized plasmids using an RNA labelling and detection kit (Roche)^87^. Probes were purified on ProbeQuant G50 Micro Columns (GE Healthcare).

Embryos were microinjected at 2-4 cell with mRNA and DNA respectively and let to develop. Embryos were fixed with 4% PFA at 60% epiboly or 24hpf. Embryos were dehydrated in 100% MeOH and dechorionated. *In situ* hybridization was carried out as previously described^14^. *In situ* embryos were imaged on a stereo microscope with uplighter in 70% glycerol. 24hpf embryos were deyolked and flatmounted under coverslips in 70% glycerol. The total length of the forebrain primordium expression of *pax6a* and negative stain of midbrain primordium before the start of hindbrain *pax6a* expression was measured in FIJI. Maximum width of hindbrain primordia was also measured in FIJI.

### Intensity of *gbx1* expression analysis

FIJI software was used to plot the pixel intensity along 3x lines spanning *gbx1* expression from the animal to vegetal pole at 60% epiboly. Expression level by pixel intensity and distance of expression domain was recorded and averaged per embryo. Data were normalised to maximum control grey value and expression distance was aligned for comparison.

### Statistical analysis

Statistical analysis carried out using IBM SPSS Statistics 26. Normal multiple comparisons were tested with one-way ANOVA. Non-normal multiple comparisons were tested using Kruskal-Wallis tests including Bonferroni correction for multiple tests if appropriate. Statistical significance: * ≤ 0.05, ** ≤ 0.01, *** ≤ 0.001.

### Computer simulations

The expansion of the neural plate was modelled *in silico* via agent-based simulation in the Chaste^43, 44^ C++ package on a HP Z840 workstation over an Intel Xeon E-series architecture. Forces between cells were defined via a Delaunay-Voronoi triangulation^88, 89^. Radially acting forces resolved along the outward pointing normal of a circle were used to represent the surface of a yolk over which the tissue grew. A similar, but inwardly directed force was used to promote intercalation and convergent expansion around the yolk. Cell division followed a renewal process with uniformly randomly drawn division times (reset following a mitotic event). Cells underwent apoptosis following a Poisson process with a pre-defined probability per unit time.

Prior to simulation, the number of cytonemes for each source cell was set following a Poisson distribution with the experimentally observed values. The tip of each cytoneme was loaded with a uniform value of Wnt. At each subsequent time step, cytonemes were either expanded or fully retracted. During growth, cytonemes grew by an amount drawn from a normal distribution with mean equal to the experimentally observed (but scaled) values. Negative sample values were discarded and redrawn. The angle of cytoneme growth was drawn from a normal distribution centred on the tangent to the yolk (so that cytonemes grew preferentially towards the receptive tissue). Cytoneme retraction followed a Poisson process with uniform probability across all cytonemes. Cytonemes exceeding a maximum length as well as cytonemes with no remaining tip Wnt were also retracted. Retracted cytonemes were replaced instantaneously with another cytoneme with zero length at the same source cell and their tip Wnt values were reset, so that the total number of cytonemes in the simulation was conserved. Cytoneme retraction and growth properties were set according to experimentally observed values where data were available or were selected so that simulations with control values were matched to control experiments.

At each time step, any cytoneme tip in contact with a receptor cell, (defined as being when the Euclidean distance between the tip and cell centre fell below the cell radius), deposited a fraction of its tip-loaded Wnt to the receptor cell in a probabilistic fashion according to a Poisson process. Following successful deposition, the Wnt values of both the cytoneme tip and receptor cell were updated. The deposition fraction was set so that control cytonemes contacted only one cell, whereas Vangl2 over-expressed cytonemes could make up to five contact events (though these could potentially be with the same cell). Wnt levels in the receptor cells decayed exponentially with constant rate between contact events.

At the end of the simulation (at t = 10h), receptor cells were marked with either a hindbrain or non-hindbrain fate according to whether or not their Wnt level was above a threshold of 100. Following this, cell fates were binned according to the cells’ angular polar coordinate around the yolk, with the origin set to be the leading edge of the source margin. Within each bin, the proportion of cells acquiring a hindbrain fate in each bin was calculated. These data were then averaged over the same bins for 100 simulations per condition. The MHB was defined to be the first bin for which this averaged proportion fell below half (so that without a bin, the majority of cells did not acquire a hindbrain cell fate). All simulation results were analysed in Python 3.6.3 (installed with Anaconda, https://www.anaconda.com) using the standard NumPy and SciPy libraries.

---

**Supplementary Figure 1:**
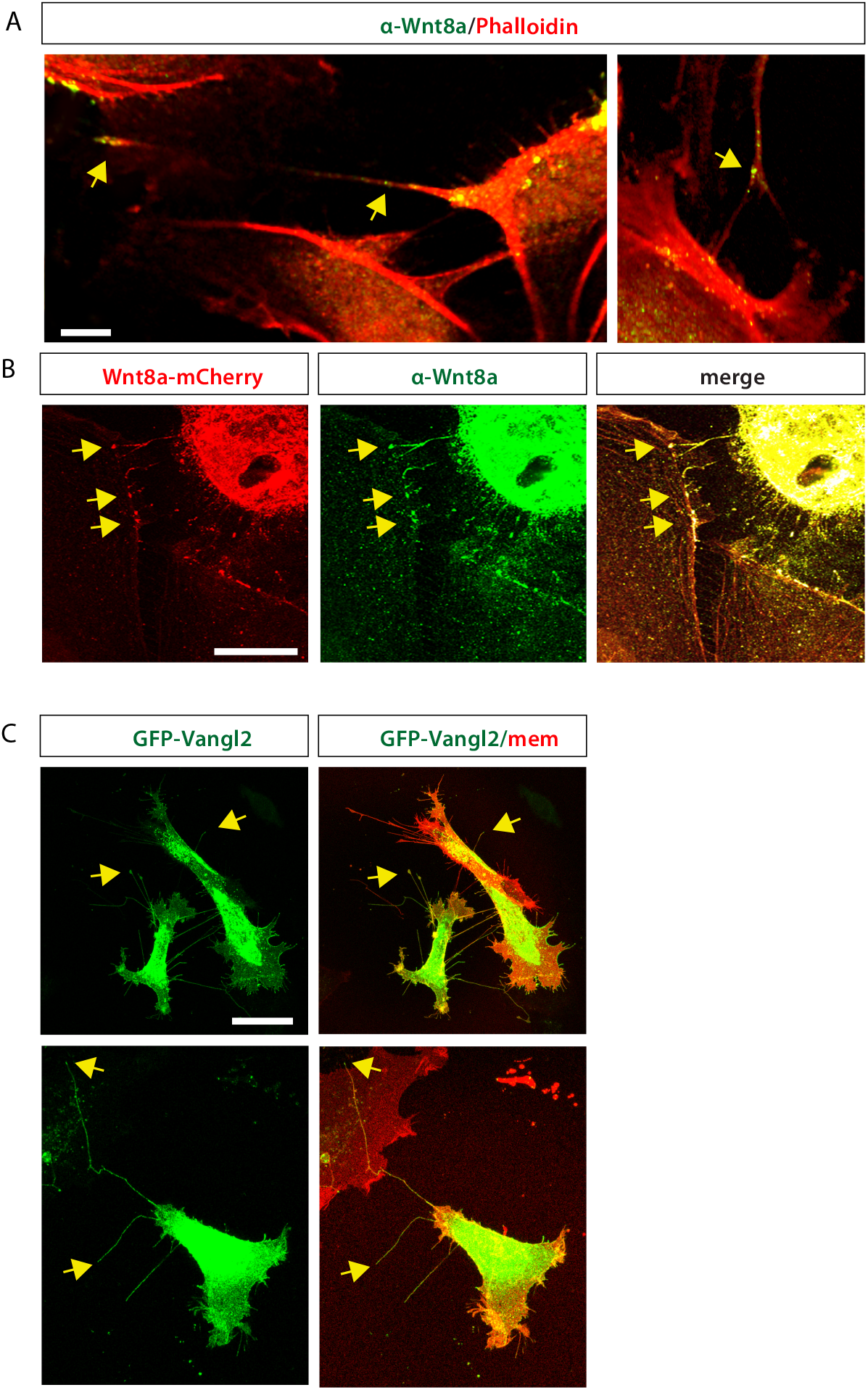
Endogenous Wnt8a is localised at cytoneme tips. (A): PAC2 cells immunostained for anti-Wnt8a with FITC-labelled secondary antibody and counterstained with Phalloidin-TRITC. Yellow arrows mark endogenous Wnt8a expression on cytoneme tips and cytoneme protrusions. Scale bar= 5μm. (B): PAC2 cells transfected with Wnt8a-mCherry and immunostained with anti-Wnt8a with FITC-labeled secondary antibody. Panels show red (Wnt8a-mCh), green (anti-Wnt8a) and merge channels. Yellow arrows show co-localisation of transfected Wnt8a-mCh with anti-Wnt8a antibody on cytoneme tips contacting recipient cells. (C): PAC2 cells transfected with mem-mCh and GFP-Vangl2. Yellow arrows show Vangl2 accumulations on cytoneme tips. Panels show merge and green (Vangl2) channel. Scale bar= 10μm.

**Supplementary Figure 2:**
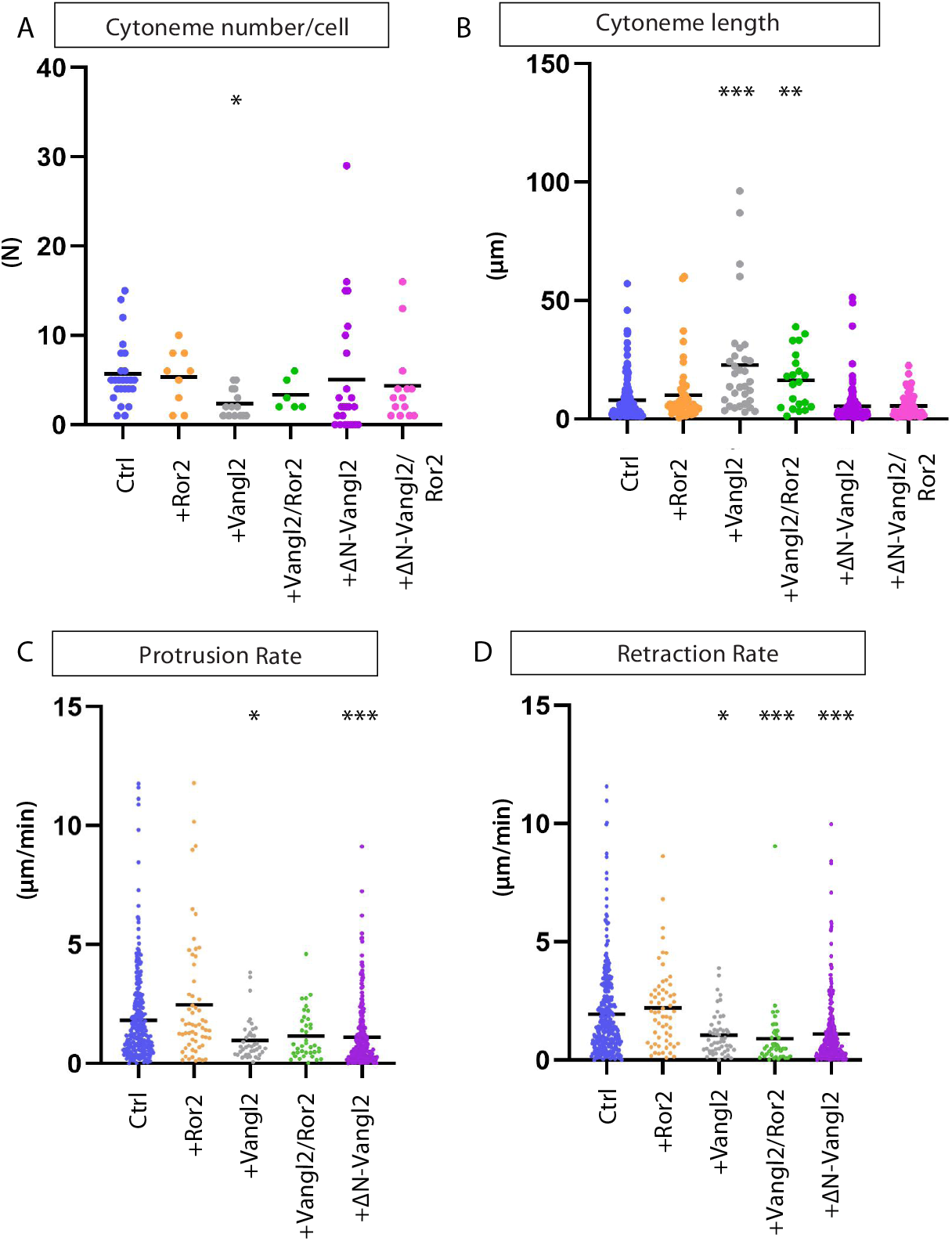
Wnt8a-positive cytoneme number, length and stability in PAC2 fibroblasts. Representations of cytoneme data from Figure 2 G, H, J, K as dot plots. (A): Number of cytonemes per cell (n= 25, 9, 14, 6, 25, 14 cells). (B): Length of Wnt8a positive cytonemes in PAC2 cells (μm). (n= 139, 52, 32, 21, 131, 65 cytonemes). (C): Protrusion rate (μm/min) of filopodia. (n=340,59, 42, 38, 327 timepoints). (D): Retraction rate (μm/min) of filopodia. (n= 340, 59, 52, 41, 327 timepoints). Statistical significance: * ≤ 0.05, ** ≤ 0.01, *** ≤ 0.001 (G,H,JK): Kruskal-Wallis tests with Bonferroni correction for multiple tests.

**Supplementary Figure 3:**
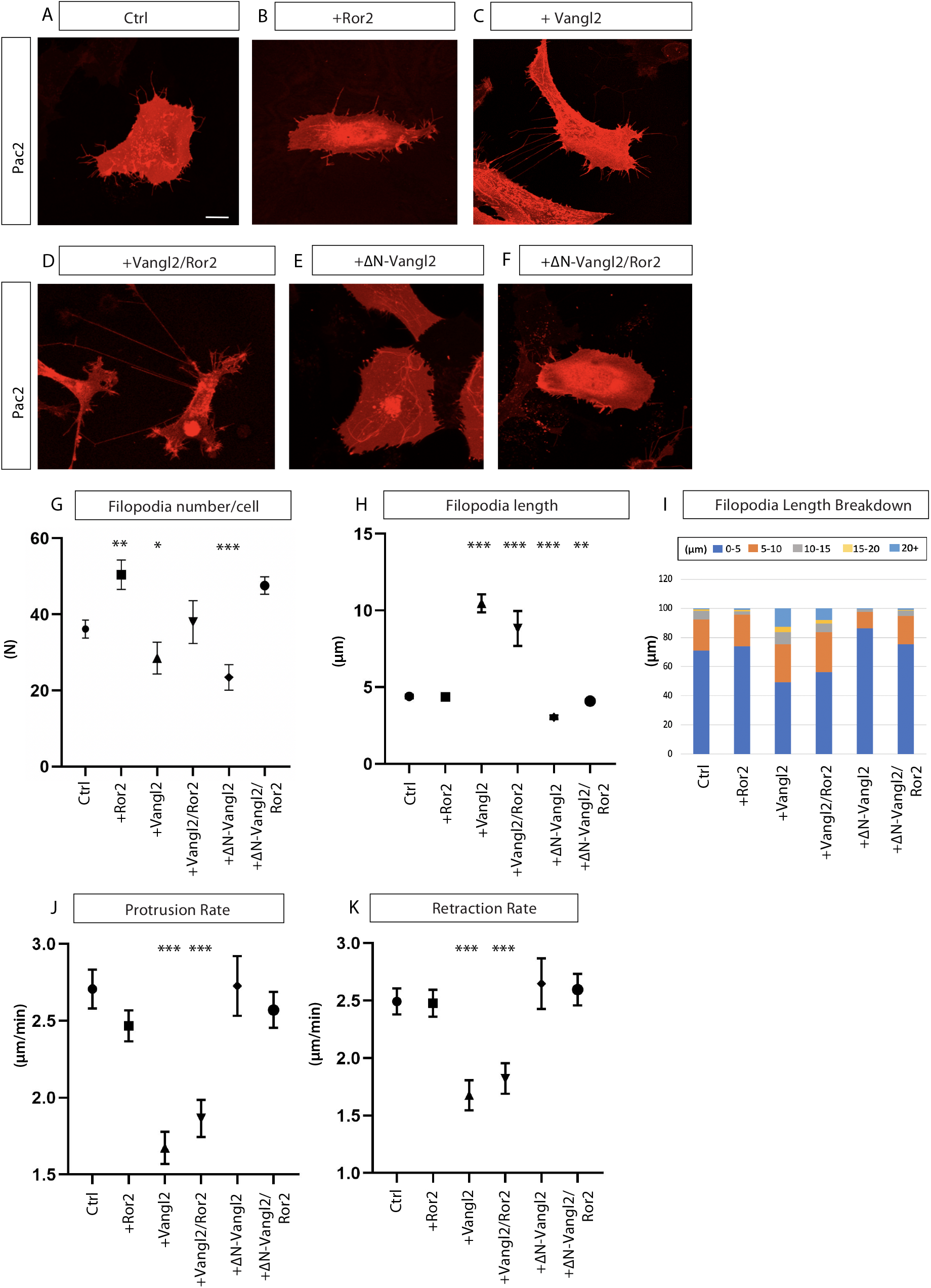
Overexpression of Vangl2 *in vitro* leads to longer, more static filopodia protrusions. (A-F): PAC2 zebrafish fibroblasts transfected with (A): Membrane-GPI-mCherry (Mem-mCh); (B): Ror2-mCherry; (C): Mem-mCh & untagged Vangl2; (D): Ror2-mCherry & GFP-Vangl2; (E): Mem-mCh & ΔN Vangl2; (F): Ror2-mCherry & ΔN Vangl2. Scale bar= 10μm. (G): Number of filopodia per cell (N = number). (n= 26, 23, 11, 5, 11, 7 cells). Stars are significant to Ror2. (H): Length of filopodia in PAC2 (μm). (n= 975, 1515, 699, 201, 423, 1371 filopodia). (I): Breakdown of the percentage of filopodia lengths into 0-5, 5-10, 10-15, 20+μm categories. (J): Protrusion rate (μm/min) of filopodia. (n= 13, 10, 10, 5, 3, 4 cells, n=209, 325, 104, 230, 119, 313 timepoints). (K): Retraction rate (μm/min) of filopodia. (n= 13, 10, 10, 5, 3, 4 cells, n= 213, 298, 90, 213, 111, 293 timepoints). Graphs represent mean and standard error of the mean. Statistical significance: * ≤ 0.05, ** ≤ 0.01, *** ≤ 0.001. ANOVA (G) & Kruskal-Wallis tests with Bonferroni correction for multiple tests (H,J,K). SEM=1. In control cells there is an average of 36.2 filopodia per cell with an average length of 4.4μm (A,G,H,L,M). 71.1% of these are short protrusions with a length below 5μm, with 21.6% between 5-10μm and 7.3% over 10μm (I). Average length of filopodia significantly increased from 4.4μm to 10.5μm in Vangl2 expressing cells (H, M, Q). This is reflected in the breakdown of protrusion lengths that were observed, with 24.4% of filopodia now within the over 10μm category and 12.7% over 20μm). Filopodia were significantly shorter in ΔN-Vangl2 and ΔN-Vangl2/Ror2 cells compared to control (E,F,H).

**Supplementary Figure 4:**
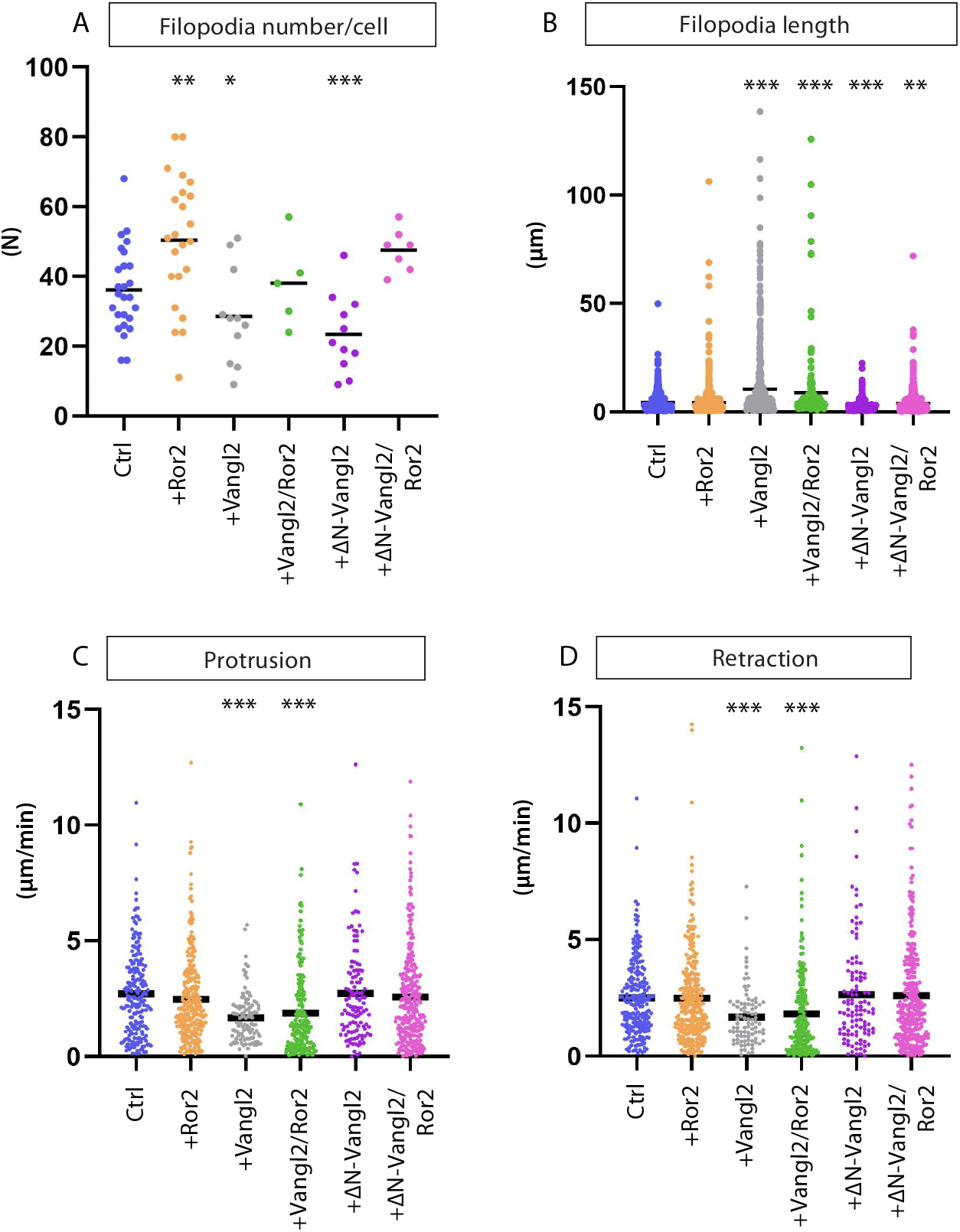
Dot plots to show overexpression of Vangl2 *in vitro* leads longer, more static filopodia. (A-D): Representations of filopodia data from supplementary figure 3 (G,H,J,K) as dot plots (A-D) respectively. (A): Number of filopodia per cell (N = number). (n= 26, 23, 11, 5, 11, 7 cells). Stars are significant to Ror2. (B): Length of filopodia in PAC2 (μm). (n= 975, 1515, 699, 201, 423, 1371 filopodia). (C): Protrusion rate (μm/min) of filopodia. (n= 13, 10, 10, 5, 3, 4 cells, n= 209, 325, 104, 230, 119, 313 timepoints). (D): Retraction rate (μm/min) of filopodia. (n= 13, 10, 10, 5, 3, 4 cells, n= 213, 298, 90, 213, 111, 293 timepoints). Statistical significance: * ≤ 0.05, ** ≤ 0.01, *** ≤ 0.001. ANOVA (A) & Kruskal-Wallis tests with Bonferroni correction for multiple tests (B-D).

**Supplementary Figure 5:**
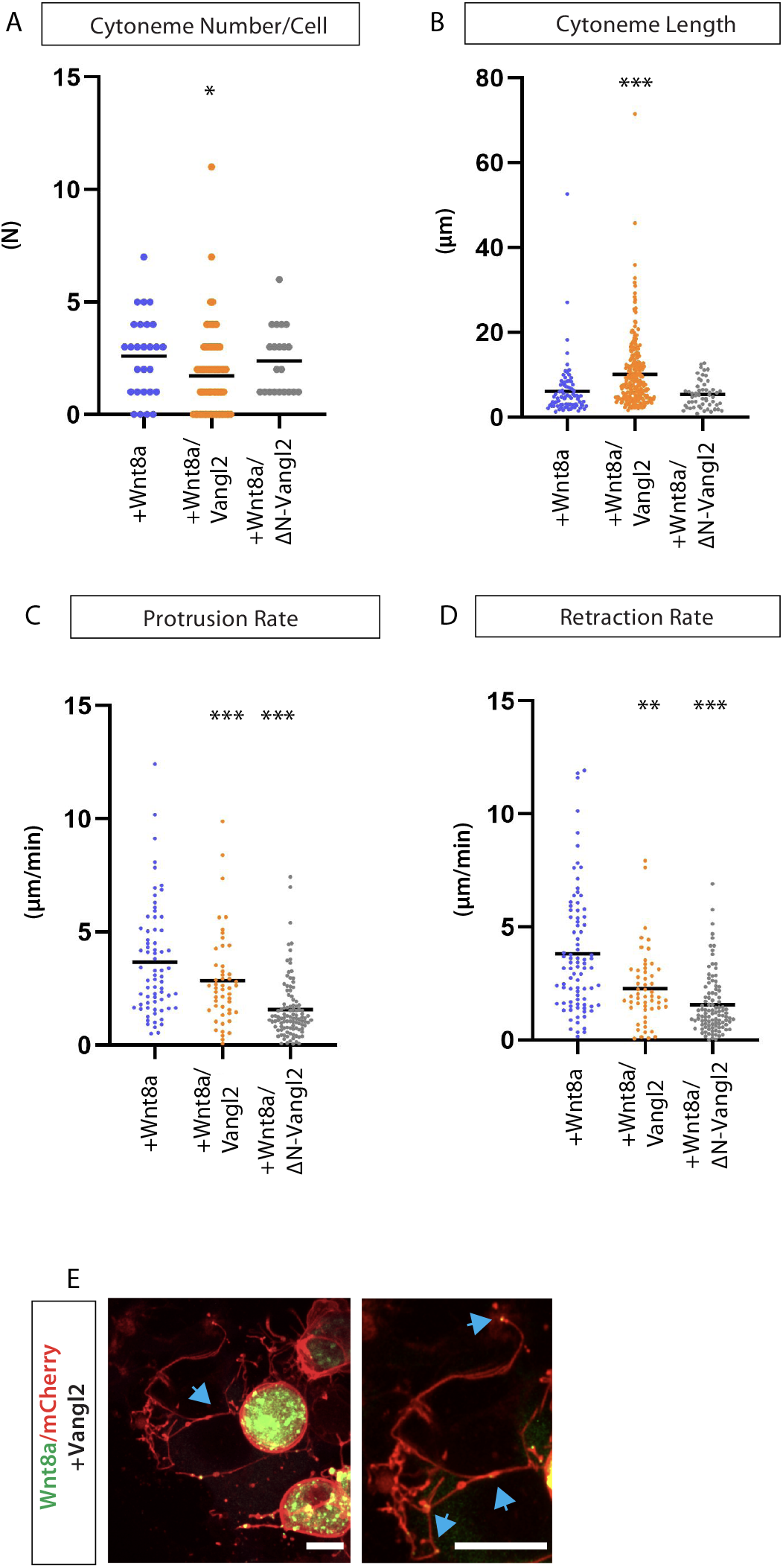
Analysis of cytoneme behaviour in zebrafish embryos. (A-D): Representations of number, length, protrusion rate and retraction rate of cytonemes *in vivo* from Figure 3 (D,E,G,H) as dot plots (A-D) respectively. (A): Number of Wnt8a positive cytonemes per cell (N = number). (n= 3, 6, 3 embryos, n= 27, 123, 21 cells). (B): Length of Wnt8a positive cytonemes (μm). (n= 81, 269, 54 cytonemes). (C): Protrusion rate (μm/min) of filopodia. (n= 3, 5, 3 embryos, n= 70, 50, 104 timepoints). (D): Retraction rate (μm/min) of filopodia. (n= 3, 5, 3 embryos, n= 86, 54, 104 timepoints). Statistical significance: * ≤ 0.05, ** ≤ 0.01, *** ≤ 0.001. Kruskal-Wallis tests with Bonferroni correction for multiple tests. (E): *In vivo* Vangl2, Wnt8-GFP and mem-mCh expressing cells exhibiting branched cytonemes. Scale bar= 10μm.

**Supplementary Figure 6:**
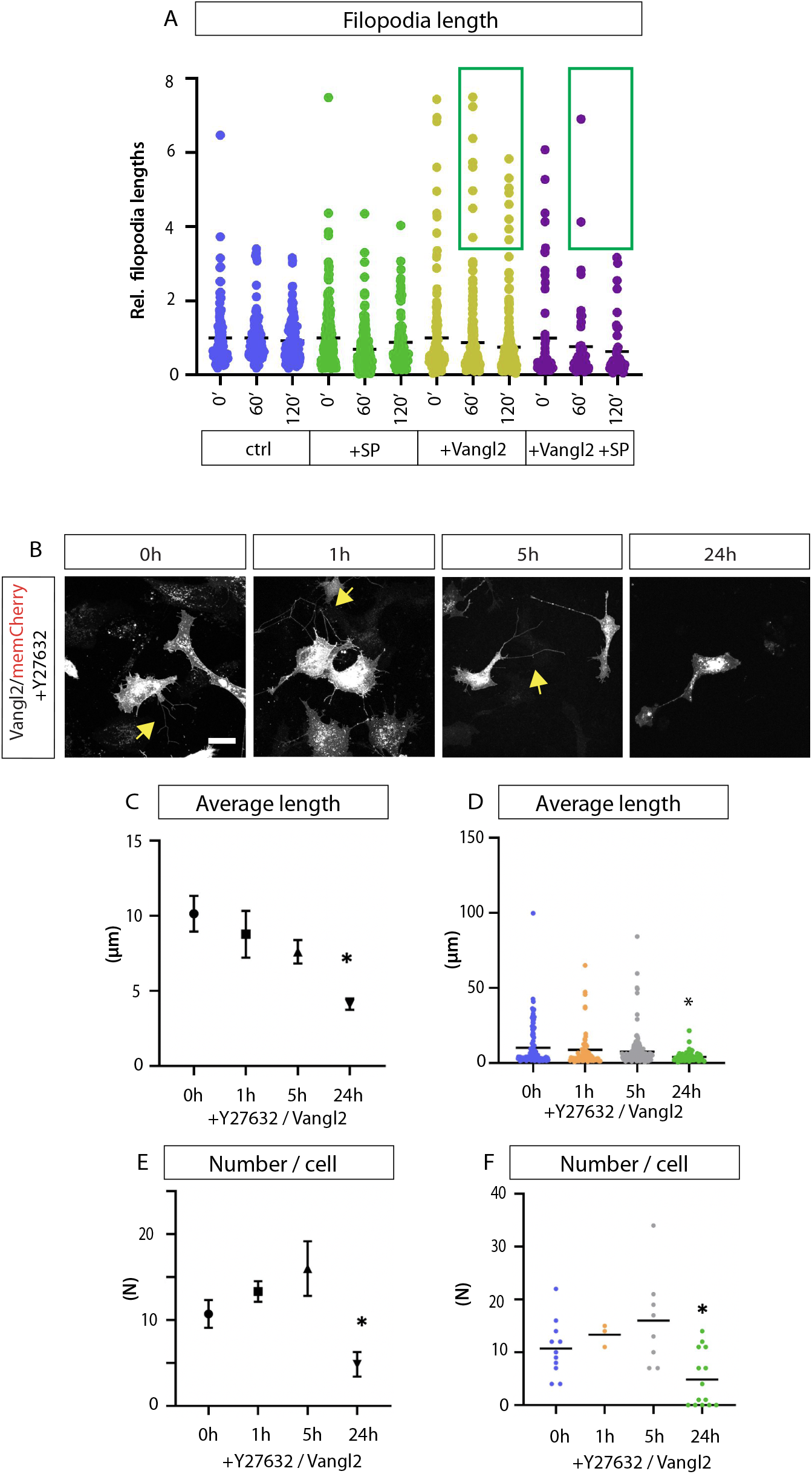
JNK and Rock inhibitor treatment on PAC2 fibroblasts. (A): Representation of Figure 4F as dot plots: Relative filopodia length (μm) after JNK inhibitor-SP600125 in relation to time=0hrs, at 0min, 60min, 120min. (n= 3, 6, 10, 4 cells. n= filopodia at 0hr, 1hr, 2hr = (102/111/109, 251/156/122, 158/185/215, 58/50/44). Green box shows the link between Vangl2/JNK signalling and long filopodia. (B): PAC2 fibroblasts transfected with GFP-Vangl2 and treated with Rock inhibitor (Y27632) for 1hr, 5hrs and 24hrs. (C-D): Length of filopodia in GFP-Vangl2 expressing cells, 1hr, 5hrs and 24hrs after Rock inhibitor - Y27632 - treatment. (n= 12, 5, 12, 9 cells, n= 121, 63, 171, 68 filopodia). (E-F): Number of filopodia in GFP-Vangl2 expressing cells, 1hr, 5hrs and 24hrs after Rock inhibitor - Y27632 - treatment, (n= 11, 3, 8, 14 cells). Represented as mean and Standard error (C,E) and as dot plots (D,F). Statistical significance: * ≤ 0.05, ** ≤ 0.01, *** ≤ 0.001. Kruskal-Wallis tests without Bonferroni correction. SEM=1. Scale bar= 10μm.

